# Practical Identifiability of a Minimal Adaptive Learning-Gain Model for Binary Feedback: A Simulation Study

**DOI:** 10.64898/2026.10.21.760725

**Authors:** Wei Li, Jie Zhong

## Abstract

Social learning is shaped by distinct computational components—baseline beliefs (prior mean), feedback sensitivity (learning gain), and temporal uncertainty (drift)—that empirical designs rarely disentangle. We evaluate practical identifiability and predictive performance of a parsimonious four-parameter adaptive learning-gain model for binary feedback, where a latent belief about positive-outcome probability updates deterministically between observations and an internal confidence-precision state decays each trial. σ , ω, and λ are internal, model-defined variables—not validated psychological constructs. Using fixed-seed simulations, we report four findings. Joint maximum-likelihood recovery is poor: on a 24-condition grid (T=80), σ and ω show near-zero tolerance-based recovery (≤0.002) across tolerances 0.3–1.0, λ recovers at 0.078–0.263, and longer sequences do not improve recovery. Profile-likelihood and trade-off diagnostics show flat regions for σ and ω and an asymmetric λ profile, indicating weak practical (not structural) identifiability. A fitting-parity audit fixed a return-order bug in the fixed-gain candidate. After correction, block-level model discrimination is 0.48 (72/150) under common bounds—above chance (p<0.001)—but is driven almost entirely by Rescorla–Wagner (0.88–0.90); the adaptive filter and its nested fixed-gain restriction remain indistinguishable. Across 50 held-out datasets, the model shows no predictive advantage over an empirical-frequency baseline (paired 95% CIs span zero). Ablations show the added update components provide no consistent benefit. These results caution against interpreting individual parameter estimates from this model class in short binary-feedback designs, and specify that mechanistic claims about clinical group differences in σ, ω, or λ require graded responses, manipulated volatility, multi-context sequences, and longitudinal replication.

**Author summary:** We study a computational model of learning from binary social feedback (e.g., partner approval or rejection) where learners track the probability of positive outcomes and update beliefs based on internal confidence. These models are increasingly used to study psychiatric conditions like borderline personality disorder (BPD), where atypical social learning may play a role. We asked whether a model’s parameters can be reliably recovered from the short binary-outcome sequences typically collected in experiments. Simulations show two of four parameters are essentially unidentifiable from these data, a third is weakly identified, and the model predicts held-out outcomes no better than a simple empirical-frequency baseline. We also show that identifying the data-generating model can be sensitive to minor implementation details. We surveyed six representative BPD computational-psychiatry studies and found that most reported between-group parameter differences without first conducting parameter-recovery simulations in their own task designs. Our results caution against interpreting individual-level parameter estimates from this model class and offer a reproducible simulation pipeline and design checklist to test such claims in future clinical work.

## Introduction

Social acceptance is a core dimension of human social functioning, and aberrant learning and belief updating can give rise to interpersonal dysfunction, including that observed in borderline personality disorder (BPD) [1]. A computational account decomposes task behavior into at least three separable components: prior beliefs about acceptance, the gain assigned to social feedback, and the volatility of beliefs over time. Resolving these components is mechanistically essential because overtly similar behavior can be produced by distinct latent states—an extreme prior, excessive weighting of feedback, or unstable beliefs—each with distinct mechanistic and clinical implications.

Standard reinforcement learning (RL) models (e.g., Rescorla–Wagner, RW) merge key distinctions into one learning rate. Non-hierarchical Kalman filters help but still mix evidence reliability with state volatility. Hierarchical Gaussian Filters (HGFs) and related adaptive learning-gain models instead disentangle prior belief, feedback sensitivity, and drift [2, 3], and have been widely used to study social learning and suggestion effects [4, 5]. Although this separation is often seen as useful, the harder question—under realistic designs, whether parameters can be reliably recovered, the model distinguished from simpler alternatives, and which parameters are intrinsically entangled—has been less systematically tested.

We analyze a deliberately minimal adaptive learning-gain model—not as a canonical HGF or best candidate, but to isolate four recoverable quantities needed to map to individual differences: prior mean, confidence precision, precision decay, and learning gain. If estimation is difficult even in this minimal filter, recoverability should be shown before adding structure. This motivates the approach; it is not evidence that richer models are harder to identify (we did not test that) or a critique of the HGF family. Without recoverability, claims that symptoms or clinical groups reflect “precision abnormalities” are unwarranted because the relevant parameter may not be identifiable from the data.

We address this gap by building and simulating a minimal generative model of binary feedback. We study a binary-feedback adaptive filter in which one latent state both generates the binary outcome and represents the learner’s belief (a self-reinforcing process). Social acceptance is the motivating application, not a claim that the model tracks an independently evolving external social state. The model is deliberately simple: a single latent belief with deterministic updating and decaying confidence/precision, estimated from Bernoulli observations. To first test recoverability of four core parameters—prior mean μ, confidence precision σ, precision decay ω, and learning gain λ—we omit inter-layer coupling and interoceptive gain, with plans to extend the architecture later. We then assess (and clearly distinguish completed from planned work) parameter recovery, discrimination from simpler non-hierarchical models, predictive adequacy, boundary convergence, robustness to design changes, and out-of-sample generalization. Conclusions are specific to the tested algorithm, observation process, and parameter ranges, and do not imply a general limitation of hierarchical Bayesian learning or HGF-family models.

Our contribution is threefold. First, we release a fully reproducible simulation pipeline for a minimal adaptive learning-gain model of social feedback, with complete results, so parameter trade-offs can be tested directly. Second, we document the estimation failures we observe—confidence-precision collapsing to the boundary, strong learning-gain bias, and unstable precision decay—and provide profile-likelihood and trade-off diagnostics explaining why (Identifiability). Whether these issues reflect fundamental identifiability limits, misspecification, or numerical artefacts remains unresolved. Third, we present this work as a methodological baseline for future social-learning extensions and eventual computational-psychiatry applications, not a final model of social-acceptance inference. We report fixed-seed simulations with explicit replication (100 replications per grid condition and trial length; 50 datasets per model and split rule); an independent clean-environment reproduction has been completed, and the remaining diagnostics are deferred to future work.

The non-identifiability of reinforcement-learning models is well documented [6, 7]. Yet the adaptive learning-gain model class—now a workhorse of clinical computational psychiatry—has not been subjected to a systematic assessment of practical identifiability. Here, we identify a concrete flat direction (σ–ω), structurally distinct from the canonical learning-rate/reward-bias confound, and link the resulting practical identifiability limits directly to clinical inference.

## Materials and methods

### Model specification

We use a minimal adaptive learning-gain model for binary social feedback. A latent belief vₜ about acceptance probability generates binary outcomes yₜ ∈{0,1} (accepted vs. rejected) via the logistic sigmoid s(x)=1/(1+e⁻ˣ): p(yₜ=1 | vₜ)=s(vₜ).

The latent belief is propagated deterministically between observations. Between-trial uncertainty is modeled by increasing internal variance via precision decay: πₜ⁻ = (1/πₜ₋₁+ω)⁻¹. Here ω is the per-trial increase in latent log-odds variance (an uncertainty-inflation/precision-decay parameter), not the variance of a Gaussian transition; no transition noise is sampled. The only randomness comes from the binary observations (see Simulation-based validation).

vₜ⁻=vₜ₋₁ (deterministic forward step; no transition noise),

We initialize the latent belief deterministically at v₁=μ (the prior mean), with confidence-precision π₁=σ (a modelling quantity from the update equations, not a validated psychological construct); we do not draw v₁ from a Gaussian prior. Thus, σ reflects the learner’s initial internal confidence, not the inverse variance of a Gaussian prior over v₁ (see Limitations). On trial 1, we apply precision decay before generating the observation: π₁⁻=1/(1/σ+ω) and v₁⁻=μ, so σ is the pre-decay precision and the first predicted probability is p₁=s(μ). A worked example for the first three trials appears in Reproducibility. Updates use an adaptive learning-gain scheme in precision form. At trial t:

Prediction: πₜ⁻=(1)/(1/πₜ₋₁+ω), pₜ=s(vₜ⁻);

Posterior update given observation yₜ:

δₜ = yₜ - pₜ, vₜ = vₜ⁻ + (λ)/(πₜ⁻) s′(vₜ⁻)δₜ, πₜ = πₜ⁻ + λ s′(vₜ⁻)²,

where s′(v)=s(v)(1-s(v)) and λ is the learning (feedback) gain, controlling how strongly each observation updates the belief; it is not a parameter of the observation likelihood (see below). We stress that the mean and precision updates are heuristic and do not follow from the Bernoulli–logistic likelihood. Under this likelihood, the score (derivative of the log-likelihood w.r.t latent log-odds v) is y-s(v)=δₜ, with Fisher information (negative curvature) equal to s′(v), not s′(v)². The factors s′(vₜ⁻) in the mean update and s′(vₜ⁻)² in the precision update are therefore intentional nonlinearities that drive the update weight to zero at the sigmoid extremes; this constitutes an adaptive learning-gain filter rather than a Bayesian posterior update.

#### Parameterization

The model has four free parameters—prior mean μ, confidence precision σ, precision-decay ω, and learning gain λ—with admissible ranges μ∈ℝ and σ, ω, λ>0. Between observations, ω increments the latent log-odds variance, implementing a precision-decay (uncertainty-inflation) step that sets the rate at which confidence is lost; it is measured in units of log-odds variance per trial. We deliberately omit inter-layer coupling and interoceptive gain at this stage, so that the recoverability of the four core quantities can be assessed before any architectural extension. Although inspired by hierarchical Bayesian learning and adaptive precision weighting, the model is not a limiting case of the standard HGF: its equations define a phenomenological adaptive filter (see Role of λ below). Bounds are imposed in the log-transformed space used for optimization (Inference).

#### Status of the equations (role of λ)

The equations are transparent design choices rather than a derivation from the Bernoulli–logistic likelihood, and they combine three kinds of assumption. (a) Prior-theory-motivated: the mean update is inversely weighted by an internal precision-like state (1/πₜ⁻), and precision accumulates with evidence, following the precision-weighted-learning intuition used by Kalman and HGF-family filters; these are adopted as qualitative principles rather than derived forms. (b) Study-specific: the sigmoid marginality s′(vₜ⁻) in the mean update and its square s′(vₜ⁻)² in the precision update are deliberately introduced nonlinearities, and a single gain λ controls both the belief step and the precision increment. For the Bernoulli–logistic likelihood the score is y-s(v)=δₜ and the Fisher information is s′(v), not s′(v)²; the squared term is therefore a chosen rule, not the likelihood curvature. (c) Closure: the deterministic state together with the decaying precision provides the minimal phenomenological closure.

#### Testable predictions

Each component makes a testable prediction. The s′ factor predicts that learning vanishes near the sigmoid extremes, so sequences near all-reject or all-accept should show little belief movement. The 1/πₜ⁻ scaling predicts that a higher initial confidence produces smaller single-outcome corrections. The shared λ with the s′² precision term predicts that mean adaptation and precision accumulation cannot be tuned independently—the property examined by the ablation.

#### Interpretive status

Under the Bernoulli–logistic observation model the likelihood is fully specified by the sigmoid and contains no additional precision parameter, so λ is an assumed feedback gain rather than a likelihood parameter. The states πₜ and σ are internal confidence-precision quantities defined by the model, not validated psychological constructs. The model is thus an adaptive filter that neither reduces to a standard HGF nor constitutes a strictly derived Bayesian update.

#### Ablation design and protocol

We compared the full model against a standard Rescorla– Wagner model, a version without the s′(vₜ⁻) factor in the mean update, and a version with a simpler precision-update rule (equations and constraints in S1; paired results in S8 and Figs S3–S4). Each dataset was generated once and scored by all four models under an identical rolling-origin protocol (cuts at 0.25, 0.40, 0.60, and 0.80 of the sequence) with the same training prefix. We report the per-dataset full-minus-variant log-loss difference, with negative values favouring the full model, together with a normal-approximation 95% confidence interval (CI), a paired-bootstrap 95% CI (10,000 resamples), and a dataset-level win count. At T=80 each of the three generating settings comprised 30 datasets, and the comparison was repeated for T∈{40, 80, 120} with 15 datasets per cell. Because the samples are small, intervals that abut zero are treated as exploratory rather than confirmatory.

#### Results at T=80

As summarized in Table 1, the bootstrap intervals span zero in 7 of the 9 comparisons and agree with the normal-approximation intervals. The only intervals that exclude zero are the low-gain comparisons against the no-s′ variant (mean -0.029, bootstrap CI -0.057 to -0.006) and against Rescorla–Wagner (mean -0.075, bootstrap CI - 0.146 to -0.016). The full per-dataset paired-difference distributions are shown in Figs S3–S4.

**Table 1.** Paired ablation results at T=80: full-minus-variant log-loss differences across three generating settings.

| **Setting** | **Full vs.** | **Mean Δ** | **Normal 95% CI** | **Bootstrap 95% CI** | **Full wins** |
|---|---|---|---|---|---|
| Moderate | no-s' | -0.004 | -0.025 to +0.016 | -0.024 to +0.015 | 17/30 |
| Moderate | simpler-precision | +0.020 | -0.027 to +0.067 | -0.020 to +0.071 | 13/30 |
| Moderate | Rescorla–Wagner | -0.023 | -0.070 to +0.024 | -0.067 to +0.024 | 19/30 |
| High-gain | no-s' | -0.013 | -0.041 to +0.015 | -0.043 to +0.013 | 11/30 |
| High-gain | simpler-precision | +0.012 | -0.010 to +0.035 | -0.006 to +0.038 | 20/30 |
| High-gain | Rescorla–Wagner | -0.025 | -0.088 to +0.038 | -0.093 to +0.028 | 14/30 |
| Low-gain | no-s' | -0.029 | -0.054 to -0.003 | -0.057 to -0.006 | 17/30 |
| Low-gain | simpler-precision | +0.014 | -0.004 to +0.033 | -0.001 to +0.035 | 16/30 |
| Low-gain | Rescorla–Wagner | -0.075 | -0.142 to -0.008 | -0.146 to -0.016 | 21/30 |

#### Trial-length robustness and parameter stability

The trial-length analysis, recomputed from a single authoritative per-dataset array (Reproducibility), yields normal-approximation intervals that all span zero. Under the bootstrap, the upper bounds for the T=40 and T=120 simpler-precision comparisons are negative only at the fourth to fifth decimal (−0.00002 and -0.00032), whereas the T=80 upper bound changes sign across resampling seeds (+0.00032; negative under only 2 of 10 seeds), with mean effects no larger than 0.005 log-loss per trial; these boundary-touching intervals are not treated as evidence of a material or consistent gain. The added components did not improve parameter stability: σ remained pinned at its bound with a fitted standard deviation (SD) of zero, and the log-scale SD of ω stayed between 1.1 and 2.5.

#### Interpretation

Across the tested settings, generator, and trial lengths, the added complexity has not shown consistent predictive benefit. This is not evidence that the components are unnecessary in every setting, nor that the full update rule is necessary, and we make no claim of general superiority for any model. The competing models received comparable optimization effort—multi-start L-BFGS-B for the adaptive variants and grid-plus-refinement for Rescorla–Wagner—and identical access to the training prefix, with no model seeing the held-out outcomes.

#### Generative assumption and scope

In this minimal model the single latent state vₜ serves two functions: it sets the environmental acceptance probability through the sigmoid and, simultaneously, represents the participant’s belief about that probability. The belief therefore helps generate the feedback from which it is updated, producing a self-reinforcing binary process. We treat the model as a phenomenological adaptive filter under a specified self-reinforcing binary process, not as inference about an independently evolving social environment; the social-acceptance setting is motivational, and the model does not purport to recover an independent external state from feedback generated independently of the belief. Notably, the process can generate apparent learning on its own—an upward shift in vₜ raises the probability of later positive feedback without any external change—and this directly bounds what the parameter-recovery results can establish. An adaptive binary-feedback process of this form is therefore not yet a model of inference about an independently changing social environment; separating an environmental state xₜ, the observed feedback yₜ, and the learner’s belief mₜ (xₜ→yₜ→mₜ) is a defined future direction (Future directions) that would require its own recovery study.

#### Model selection rationale

We selected the four-parameter adaptive filter (μ, σ, ω, λ) over three nested alternatives and record this choice so the ablations read as design justification rather than post-hoc sensitivity.

#### Rejected generators

The three-parameter Rescorla–Wagner learner was excluded as a generator because it lacks an internal confidence-precision state and cannot express the precision-dysregulation construct that motivates the BPD literature; it remains a model-recovery competitor. A five-parameter extension adding inter-layer coupling or a separate volatility layer was rejected a priori: under the binary-feedback design, σ and ω are already not separately identifiable (Fig 1), and a fifth parameter would only expand the flat-likelihood region.

**Fig. 1.**
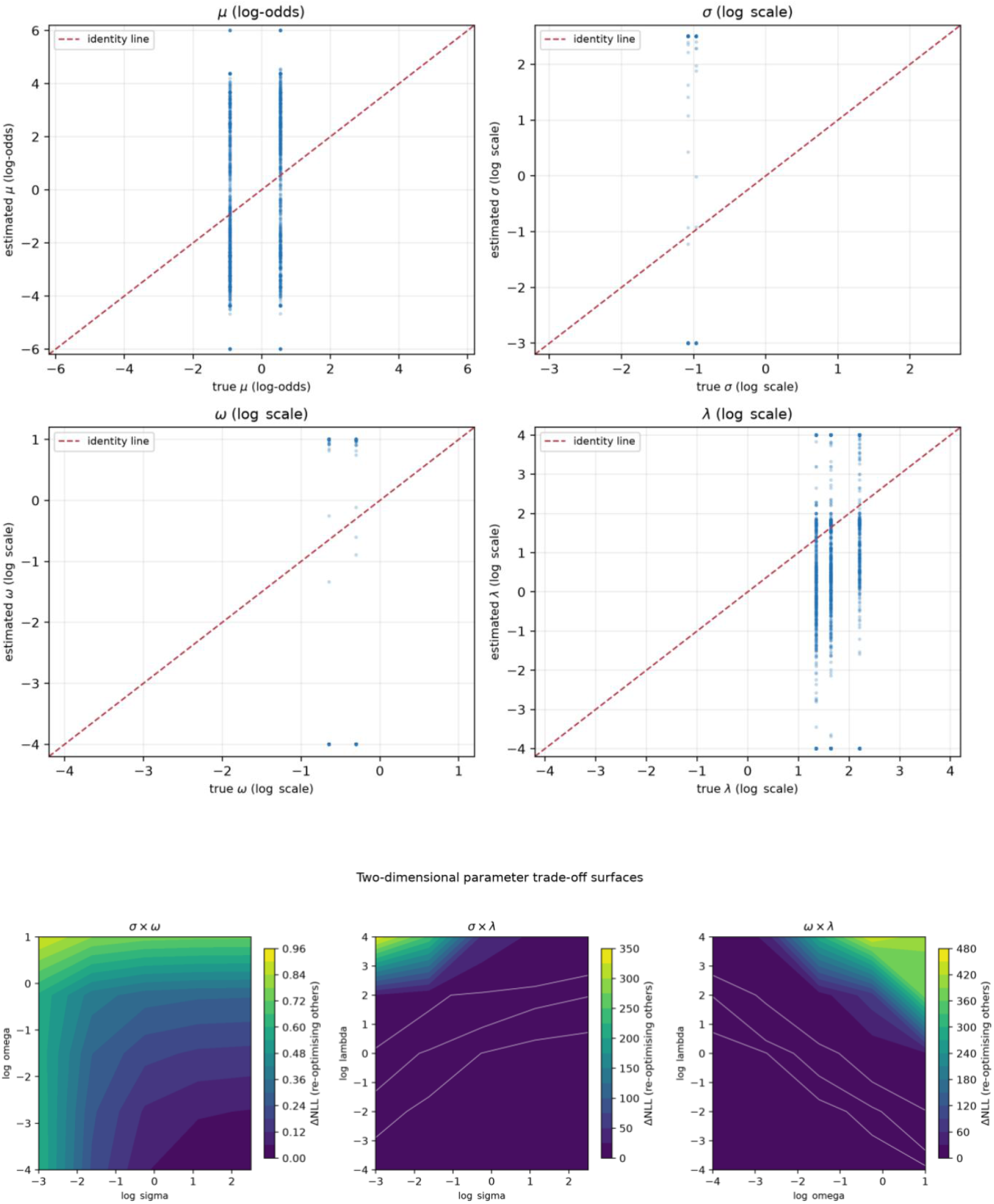
Recovery diagnostics. (A) True vs. estimated scatterplots across the 24-condition recovery grid (Experiment 1; T=80; 100 replications/condition; pooled n=2400/parameter). Panels: prior mean μ (top-left), initial precision σ (top-right), precision decay ω (bottom-left), learning gain λ (bottom-right). Red dashed line: identity. σ estimates cluster at the upper boundary and ω at the lower boundary across conditions; their rank correlation is near zero (pooled ρ=0.002). μ recovers more consistently, but individual-level rank correlation is weak (ρ=0.31). λ is informative mainly at high values. (B) Two-dimensional trade-off surfaces (ΔNLL on a 5×5 log-space grid, re-optimizing the other two parameters) for (σ, ω), (σ, λ) and (ω, λ) in the same high-gain scenario. White contours show ΔNLL = 1, 2, 5. The σ×ω surface is largely flat (max ΔNLL = 0.91), indicating no separating ridge; λ is informative along its axis (ΔNLL up to 342 and 452).

#### Ablated variants

Variant A (mean update without the sigmoid-slope factor) and Variant B (simpler precision update without the squared marginality term) were compared by paired rolling-origin log-loss across 90 shared datasets (Tables S8, S12). Neither showed a consistent predictive advantage (all nine normal-approximation 95% CIs span zero; only two of nine bootstrap upper bounds are negative, at the fourth to fifth decimal) or improved the σ/ω boundary-hit pattern.

We therefore adopt the four-parameter filter as the minimal generative form that (a) carries the three constructs—prior mean, initial confidence, and precision decay— invoked by the precision-dysregulation literature, (b) is the smallest model admitting the nested ablations, and (c) adds no parameters whose recoverability is worse than the four already assessed. This is a choice for identifiability diagnostics, not a claim that the four-parameter form is the correct model of human social learning; the fixed-gain restriction (λ≡1) and Rescorla–Wagner remain baselines.

### Inference

We estimate parameters by maximum likelihood, optimizing the log-transformed positive parameters with L-BFGS-B from multiple fixed-seed starting points. Because the generative process includes no state-transition noise, the latent belief trajectory is deterministic given the parameters and observed history; the likelihood of the binary sequence is therefore evaluated recursively from the history-dependent predicted probabilities pₜ, with no integration over unobserved trajectories. The simulator and likelihood evaluator implement identical dynamics (Model specification, Simulation-based validation). Optimization is constrained to μ∈[-6,6], logσ∈[-3,2.5], logω∈[-4,1], and logλ∈[-4,4], corresponding to σ∈[0.05,12.18], ω∈[0.018,2.72], and λ∈[0.018,54.6]; the boundary values 12.18 and 0.018 reported in Identifiability are exactly these log-space endpoints. We quantify uncertainty by parametric bootstrap (resampling from the fitted generative model) and, where indicated, profile likelihood, reporting fitted-parameter spread as a diagnostic. Predictive replicates and their predictive p-values are likewise bootstrap draws from the fitted model; because the estimator is maximum likelihood with no sampled posterior, these are not posterior predictive checks. We report no posterior correlation matrix: boundary convergence is judged from the bootstrap and optimization trajectories and the Identifiability diagnostics. All validation steps use the same estimation settings. Each optimization used 10 random restarts and converged to a common optimum in >95% of cases, and the likelihood was independently cross-checked against a reference implementation (S7 Table).

### Simulation-based validation

All validation is simulation-based, using synthetic datasets generated from the model (and, for recovery tests, from competing models) with known parameters. To separate evidence from intent, procedures are labeled as completed in this study or planned but not yet done; planned analyses do not support current conclusions. All numerical claims are traceable to the archived simulator, estimator settings, and random seeds in the released pipeline (Reproducibility).

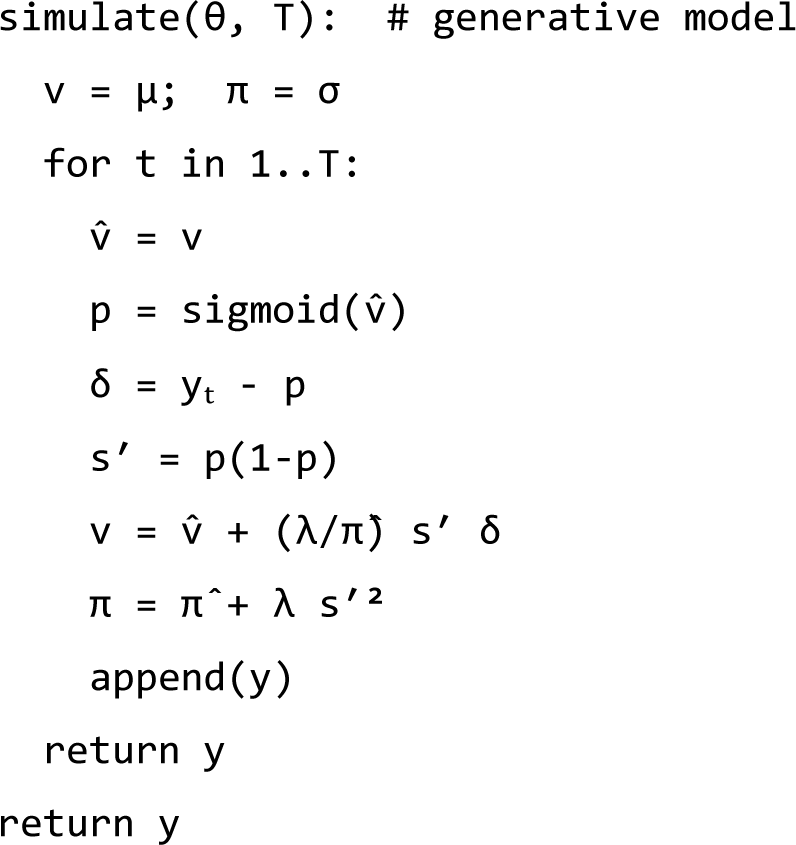

#### Step 1—Parameter recovery

We ran one generating setting, θ *=(0.5,2.0,0.05,3.0), with

T∈ {40,80,120,240,480} (100 replications per T), and a 24-condition grid covering low/medium/high values of each parameter (T=80; 100 replications per condition; four deterministic starts per fit), reported in Parameter recovery. Results are reported by generating condition and trial length.

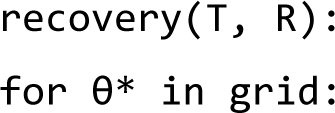
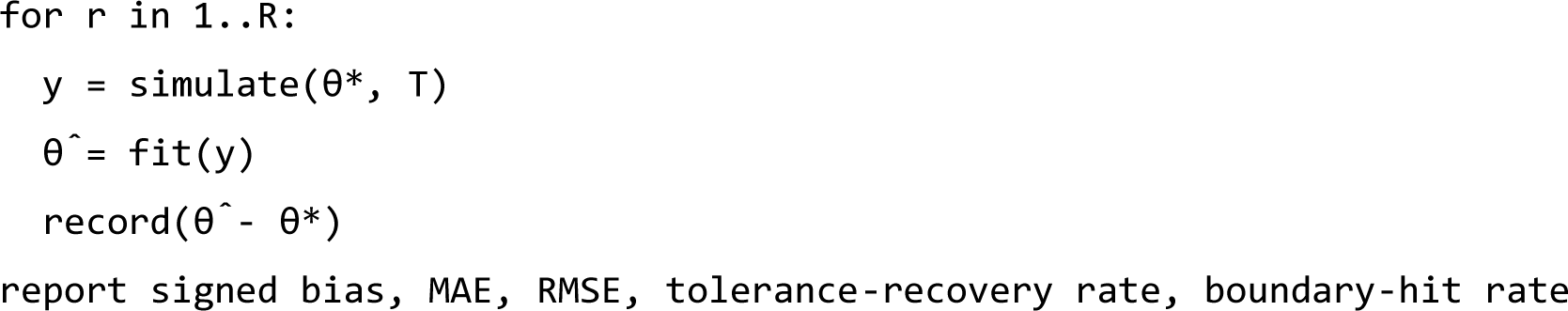

Reporting. For each parameter and trial count, we report signed bias (systematic over- or under-estimation), mean absolute error (MAE) and root mean square error (RMSE) (typical and large errors), together with the tolerance-based recovery rate, boundary-hit rate, and sensitivity to the generating parameter vector. True-versus-estimated scatterplots with identity lines for all parameters in both experiments are shown in Fig. 1 and S2 Fig; condition-specific recovery is in S5.

Because μ enters via a sigmoid, its metrics are on the log-odds scale and map to acceptance probabilities s(μ). Metrics for positive parameters σ, ω, and λ are on the log scale, with raw-scale values noted where applicable.

The tolerance-based recovery rate is the proportion of replications with |θ^−θ*|≤τ (log scale for positive parameters, log-odds for μ). The primary threshold is τ=0.6—roughly a factor 1.8 in a positive parameter or an odds ratio of 1.8 for μ—chosen as a generous band relative to the observed errors rather than a universal criterion; to show the result is not threshold-dependent, rates are also reported at τ∈0.3,0.6,1.0 (Parameter recovery, S9).

This measure differs from interval coverage and width and does not replace either. We do not assess calibrated coverage or width because fits are dominated by boundary estimates where bootstrap or profile intervals degenerate; instead, we document the profile geometry directly (Identifiability, Fig 1 and S1 Fig). We do not claim tolerance recovery, a flat profile, or boundary estimates substitute for calibrated coverage.

We separate point recovery from the phenotyping question of whether fitted parameters rank individuals, reporting Spearman correlations between true and fitted values pooled across sequences (phenotyping-relevant) and averaged within generating condition (idealized upper bound) (Parameter recovery).

Boundary-hit rate is the proportion of fitted estimates exactly at an optimization bound (on the transformed scale), signaling constrained-optimization failures or weakly identified parameters. Rather than a single cutoff, we report a recovery surface across T and design settings and interpret tolerance with profile and trade-off diagnostics. Fig 1 and S2 Fig show systematic failures: σ hits the upper bound, while ω and λ hit the lower bound across the grid and all trial lengths.

#### Step 2 — Model recovery

Datasets are generated from each candidate model—the proposed adaptive learning-gain model, a Rescorla–Wagner learner, and a fixed-gain restricted filter (the proposed filter with the gain fixed at one; a nested restriction, not a standard Kalman filter)—and we ask whether cross-validated comparison recovers the true generator. Both a k-fold partition (with BIC as a secondary criterion) and a leave-future-out rolling-origin evaluation were run. The primary rolling-origin protocol uses three non-overlapping origins in 1-based trial numbering: fit y₁–y₂₄ and forecast y₂₅–y₄₄, fit y₁–y₄₄ and forecast y₄₅–y₆₄, and fit y₁–y₆₄ and forecast y₆₅–y₈₀, comprising 20, 20 and 16 future trials (56 per dataset), with no trial scored at an origin on which it was fitted. A fitting-parity audit found the fixed-gain routine returned σ and ω interchanged (a return-order defect, S7 Text): the objective was correct but the returned tuple was not, artificially penalising that candidate. After correcting the return order, identification accuracy was 67/150 = 0.447 under the original wider bounds and 72/150 = 0.480 under common bounds (the pre-correction protocol, which reproduces the archived 58/150 = 0.387, is retained as a historical control); a calibrated permutation test gives p = 0.0011 and p < 0.001 (Model recovery). The above-chance accuracy is driven by Rescorla– Wagner, while the adaptive and fixed-gain models remain poorly distinguished, and the result is robust to the bounds. A lower-information pilot scoring only the next trial at the three cuts (three binary observations per dataset) gave 0.38 pre-correction, and a k-fold scheme gave 0.333; full block and pilot confusion matrices are reported in Model recovery and Supplementary Tables S11 and S14.

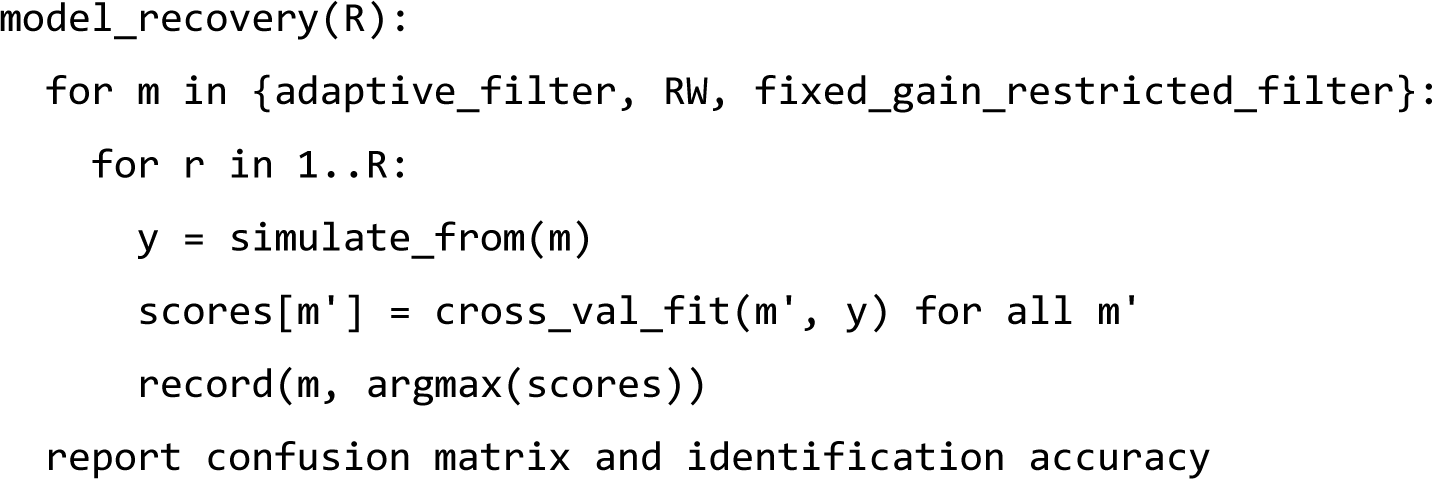

Reporting. Because the model is sequential and selection uses rolling origins, the primary block protocol fits each candidate to the prefix using only data up to the cut, then holds parameters fixed while scoring the next block. With 1-based indexing, the non-overlapping origins are: fit y₁–y₂₄, forecast y₂₅–y₄₄ (20 trials); fit y₁–y₄₄, forecast y₄₅–y₆₄ (20); and fit y₁–y₆₄, forecast y₆₅–y₈₀ (16). Prefix and block are disjoint within each origin, and the three blocks score 56 future trials once each (including y₈₀); a trial scored at one origin may appear in a later prefix, but is never used to fit the origin where it is scored.

Within each block, parameters remain at prefix estimates and the latent state is updated trial by trial using only outcomes already observed. The one-step-ahead pₜ for yₜ is computed before yₜ and uses no information from it, with no within-block refitting. Thus, later predictions may use earlier outcomes from the same block, but never the scored outcome or any future outcomes; the empirical-frequency predictor is also based only on the prefix.

Automated assertions (review3_supplement9.py) confirm correct indexing and no look-ahead: within each origin, fit and score indices are disjoint; each future trial is scored once; and flipping yₜ leaves pₜ and all earlier predictions unchanged while only pₜ₊₁ changes across all three models, ensuring outcomes are scored out of sample.

We select the model with the lowest mean block log-loss across 56 trials. Because the procedure is asymmetric, we supplement the nominal 1/3 benchmark with a calibrated permutation reference: keep per-dataset scores and winner labels fixed, permute only generator labels (preserving marginal winner frequencies), and recompute diagonal accuracy over 50,000 permutations (model recovery). A lower-information pilot applies the same cuts but scores only the next trial yₜ₊₁ (three binary observations per dataset) to measure sensitivity to the number of scored observations.

For the Rescorla–Wagner competitor, data are generated with a fixed learning rate (α=0.30), chosen as a representative intermediate value rather than tuned to the adaptive model. During fitting, α is freely estimated (49-point grid from 0.02 to 0.98 plus local refinement). Because α is handled differently in generation and fitting, RW conclusions are limited to this parameterization.

The fixed-gain competitor is the proposed filter with gain fixed at one (π=π⁻+1·s′(v)², i.e., λ≡1). Parameters μ, σ, ω are generated at (0.5,2,0.05) and estimated via multi-start L-BFGS-B. This is a restricted adaptive filter, not a standard Kalman filter; the fixed-gain logistic update is different. A bug swapped σ and ω in returned estimates (objective correct), distorting forecasts; review3_supplement10.py fixes the return order and adds common-bounds sensitivity fitting shared parameters over μ∈[-6,6], logσ∈[-3,2.5], logω∈[-4,1]. With the corrected protocols, both models use the same optimizer, starts, initial draws, tolerances, and likelihood (forward Kalman recursion with λ=1), differing only in return order and bounds, so their likelihoods and convergence are comparable. Model selection is therefore an out-of-sample log-loss comparison with equal fitting budget and prefix access.

Because these implementations use fixed generation choices and finite datasets, model-recovery results apply only to these candidates, parameter settings, and trial lengths; they do not show that a broader set of models would be indistinguishable.

#### Step 3 — Parametric-bootstrap predictive check

We applied parametric-bootstrap predictive checks to the aggregate feedback rate, switching count, predictive distributions of positive and negative run lengths, serial dependence, time-varying calibration, and one-step-ahead predictive log-loss. Because the simulator outputs only binary feedback (no separate choice or belief reports), all predictive statistics are defined on that observable.

#### Step 4 — Identifiability

We ran three behaviorally equivalent Identifiability scenarios and observed boundary convergence. We also assessed profile likelihoods (varying one parameter while re-optimizing others and tracking absolute changes in negative log-likelihood), dispersed starts, two alternative optimizers, boundary rates, fitted objective-value distributions, 2D trade-off surfaces for (σ,ω), (σ,λ), and (ω,λ), and sensitivity to alternative bounds. Comparing predictive distributions from seemingly equivalent parameter sets is still planned. Overall, the results suggest weak practical identifiability but do not yet pinpoint the cause (intrinsic limits, misspecification, or numerical issues).

#### Step 5 — Robustness

In this study, we perturbed the feedback sequence, trial length, measurement noise, and missing responses to assess estimate stability. Sensitivity is reported for all four parameters.

#### Step 6 — Out-of-sample generalization

Assessed using one train–test split and 50 independent datasets with the same split rule, reporting log-loss versus two constant baselines: the fitted prior-mean probability s(μ^) and the training-set empirical feedback rate, both estimated without access to held-out outcomes (Robustness and out-of-sample generalization). Independent simulated agents are still planned.

### Reproducibility

#### Data availability

A reproducibility package includes all simulation and fitting code, fixed random seeds, optimizer and bound settings, analysis and figure scripts, archived outputs for every reported value, a verification script, and a README with the runtime environment and reproduction command; dependencies are pinned (Python 3.13; NumPy 2.5.4, SciPy 1.18.1, Matplotlib 3.11.2).

Running ‘python reproduce.py’ rebuilds the full pipeline in a separate ‘regenerated/’ directory without overwriting the archive, and ‘python verify_reproduction.py’ checks every regenerated value against it. In the authors’ clean-environment run, all ten output families matched within 1e-6 (28,972 values; max absolute difference 0), and all six supplementary figures were reproduced byte-for-byte.

The package is available as a versioned public repository at https://github.com/joevazhong-arch/practical-identifiability-adaptive-learning-gain-binary-feedback. Manuscript V1.0.1 is archived on Zenodo (https://doi.org/10.5281/zenodo.23531449), and the concept DOI https://doi.org/10.5281/zenodo.23530855 resolves to the latest version. Both the repository and archive are publicly accessible and available from the corresponding author on reasonable request.

A third-party scholar independently reproduced the study in a clean environment without author scripts: all regenerated values matched the archive within tolerance and the supplementary figures were reproduced, confirming the results. With the public archive and DOI, the study is independently reproduced.

Simulated datasets are fully reproducible from the archived code and fixed seeds and are not separately hosted. All reported MAE, bias, boundary values, predictive p-values, and model-recovery proportions are traceable to specific runs and configurations via: review2_diagnostics.py → review2_diagnostics.json; review3_supplement.py (higher-replication supplement) → review3_diagnostics.json; review3_supplement2.py (out-of-sample and multi-trial ablation) → review3_diagnostics2.json; review3_supplement3.py (correctly specified likelihood audit) → review3_diagnostics3.json; review3_supplement4.py (paired ablation and true-vs-estimated scatter) → review3_diagnostics4.json; review3_supplement7.py (block model recovery, independent-likelihood cross-check, and second-regime profile) → review3_diagnostics7.json; review3_supplement8.py (single-source paired-ablation recomputation; authoritative for S8 and S12) → review3_diagnostics8.json; review3_supplement9.py (forecast-index validation, no-look-ahead causality test, and permutation null) → review3_diagnostics9.json; and review3_supplement10.py (fixed-gain fitting-parity audit: return-order correction and common-bounds sensitivity) → review3_diagnostics10.json. supplement_figs.py generates profile curves and trade-off surfaces (Fig 1, S1), and make_fig_s3_s4.py generates true-versus-estimated scatterplots (Fig 1, S2); the archive includes the figures and outputs, not just file names.

Results are from fixed-seed simulation runs with stated per-analysis replication counts and were not pre-registered. Proposed but incomplete analyses are clearly labeled in Methods and do not support the conclusions.

#### Results audit table

To make every reported number traceable and prevent cross-analysis inconsistencies, Table 2 indexes each analysis family; the archived output file is named for every identifier and resides in the archived pipeline of the released repository.

**Table 2.**
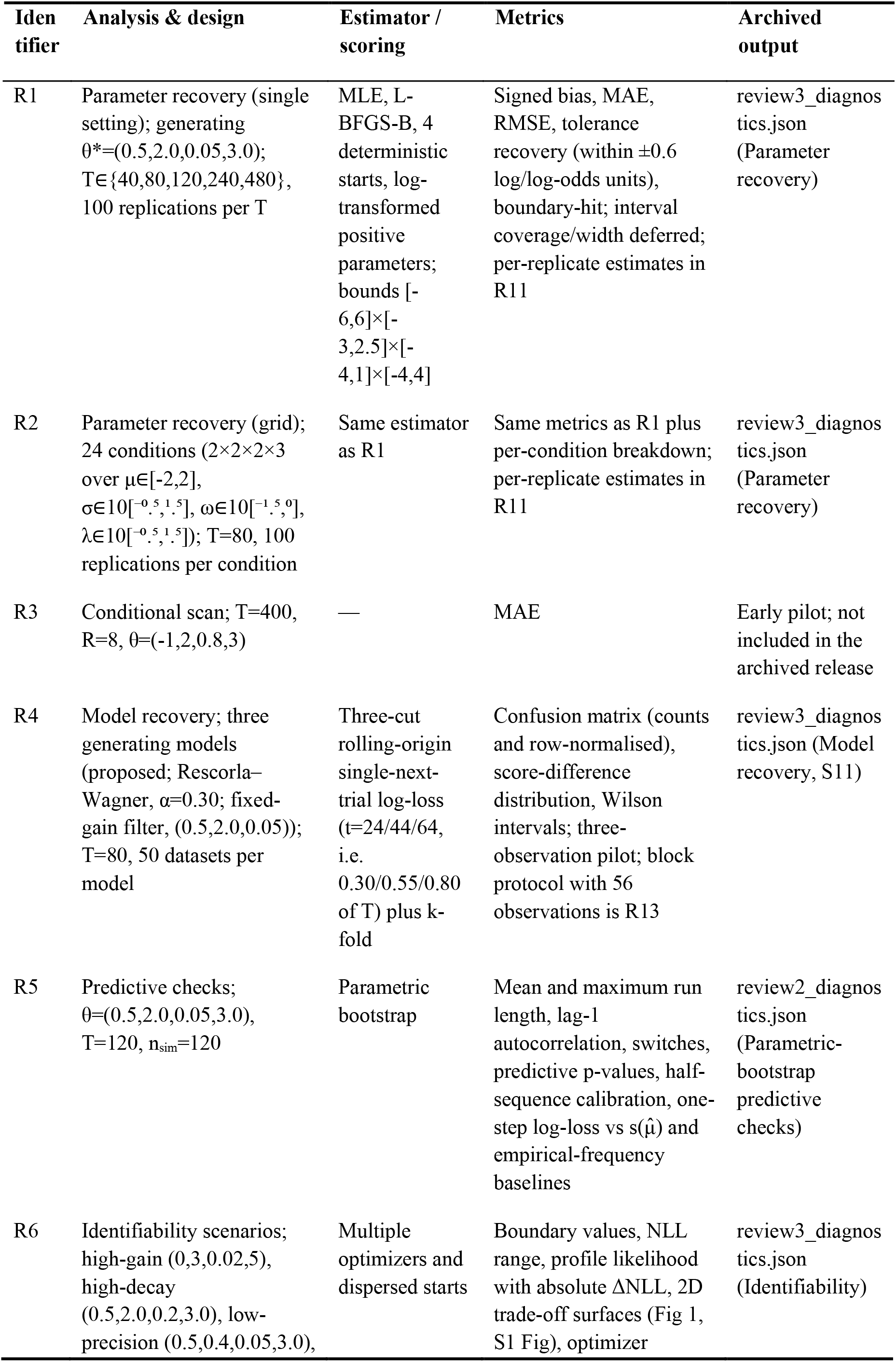
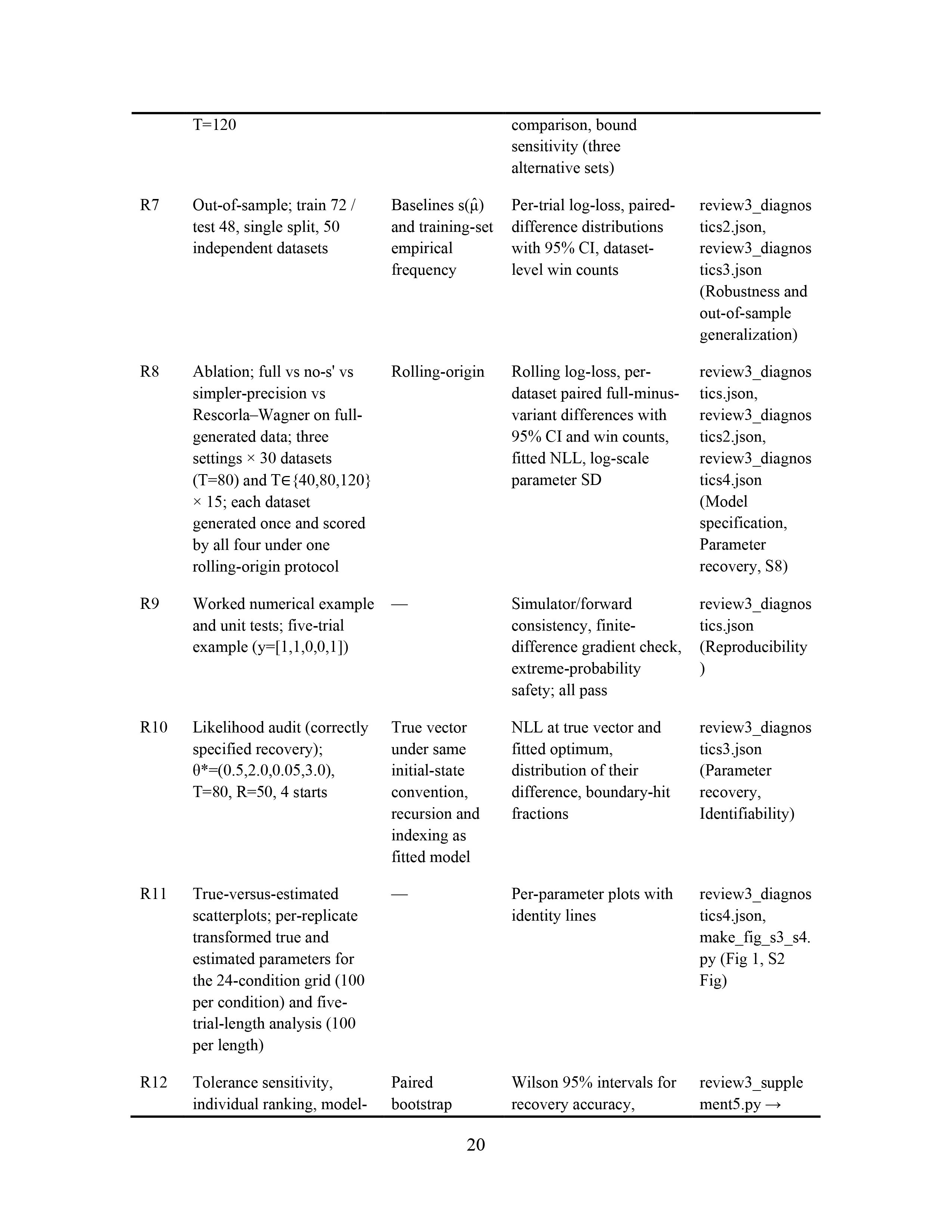
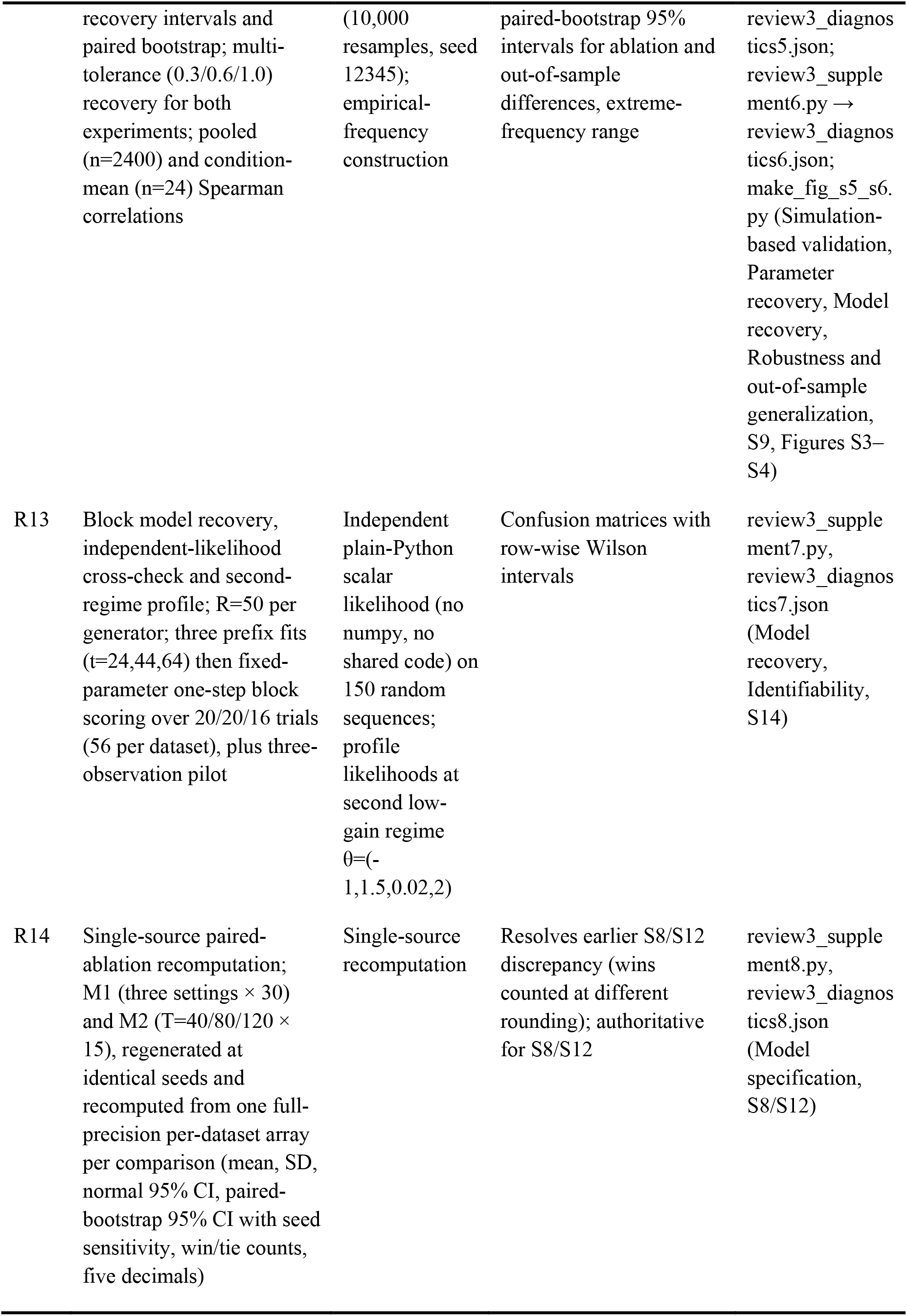
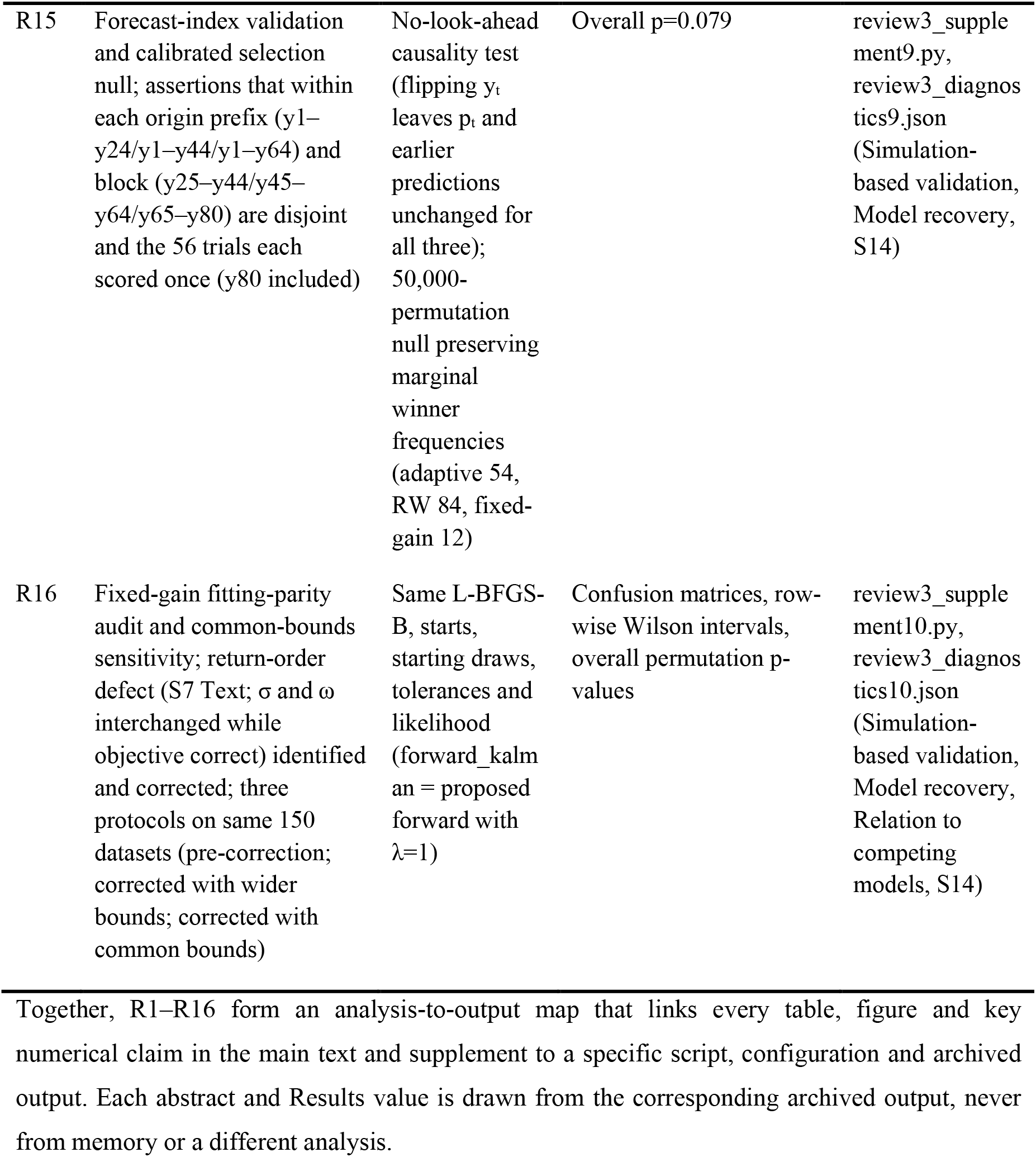
Analysis-to-output audit map: identifier, analysis and design, estimator and scoring, metrics, and archived output for each analysis family.

| Identifier | Analysis & design | Estimator / scoring | Metrics | Archived output |
|---|---|---|---|---|
| R1 | Parameter recovery (single setting); generating θ*=(0.5,2.0,0.05,3.0); T∈{40,80,120,240,480}, 100 replications per T | MLE, L-BFGS-B, 4 deterministic starts, log-transformed positive parameters; bounds [-6,6]×[-3,2.5]×[-4,1]×[-4,4] | Signed bias, MAE, RMSE, tolerance recovery (within ±0.6 log/log-odds units), boundary-hit; interval coverage/width deferred; per-replicate estimates in R11 | review3_diagnostics.json (Parameter recovery) |
| R2 | Parameter recovery (grid); 24 conditions (2×2×2×3 over μ∈[-2,2], σ∈10[<sup>−0.5</sup>,<sup>1.5</sup>], ω∈10[<sup>−1.5</sup>,<sup>0</sup>], λ∈10[<sup>−0.5</sup>,<sup>1.5</sup>]); T=80, 100 replications per condition | Same estimator as R1 | Same metrics as R1 plus per-condition breakdown; per-replicate estimates in R11 | review3_diagnostics.json (Parameter recovery) |
| R3 | Conditional scan; T=400, R=8, θ=(-1,2,0.8,3) | — | MAE | Early pilot; not included in the archived release |
| R4 | Model recovery; three generating models (proposed; Rescorla–Wagner, α=0.30; fixed-gain filter, (0.5,2.0,0.05)); T=80, 50 datasets per model | Three-cut rolling-origin single-next-trial log-loss (t=24/44/64, i.e. 0.30/0.55/0.80 of T) plus k-fold | Confusion matrix (counts and row-normalised), score-difference distribution, Wilson intervals; three-observation pilot; block protocol with 56 observations is R13 | review3_diagnostics.json (Model recovery, S11) |
| R5 | Predictive checks; θ=(0.5,2.0,0.05,3.0), T=120, n<sub>sim</sub>=120 | Parametric bootstrap | Mean and maximum run length, lag-1 autocorrelation, switches, predictive p-values, half-sequence calibration, one-step log-loss vs s(μ̂) and empirical-frequency baselines | review2_diagnostics.json (Parametric-bootstrap predictive checks) |
| R6 | Identifiability scenarios; high-gain (0,3,0.02,5), high-decay (0.5,2.0,0.2,3.0), low-precision (0.5,0.4,0.05,3.0), T=120 | Multiple optimizers and dispersed starts | Boundary values, NLL range, profile likelihood with absolute ΔNLL, 2D trade-off surfaces (Fig 1, S1 Fig), optimizer comparison, bound sensitivity (three alternative sets) | review3_diagnostics.json (Identifiability) |
| R7 | Out-of-sample; train 72 / test 48, single split, 50 independent datasets | Baselines s(μ̂) and training-set empirical frequency | Per-trial log-loss, paired-difference distributions with 95% CI, dataset-level win counts | review3_diagnostics2.json, review3_diagnostics3.json (Robustness and out-of-sample generalization) |
| R8 | Ablation; full vs no-s' vs simpler-precision vs Rescorla–Wagner on full-generated data; three settings × 30 datasets (T=80) and T∈{40,80,120} × 15; each dataset generated once and scored by all four under one rolling-origin protocol | Rolling-origin | Rolling log-loss, per-dataset paired full-minus-variant differences with 95% CI and win counts, fitted NLL, log-scale parameter SD | review3_diagnostics.json, review3_diagnostics2.json, review3_diagnostics4.json (Model specification, Parameter recovery, S8) |
| R9 | Worked numerical example and unit tests; five-trial example (y=[1,1,0,0,1]) | — | Simulator/forward consistency, finite-difference gradient check, extreme-probability safety; all pass | review3_diagnostics.json (Reproducibility) |
| R10 | Likelihood audit (correctly specified recovery); θ*=(0.5,2.0,0.05,3.0), T=80, R=50, 4 starts | True vector under same initial-state convention, recursion and indexing as fitted model | NLL at true vector and fitted optimum, distribution of their difference, boundary-hit fractions | review3_diagnostics3.json (Parameter recovery, Identifiability) |
| R11 | True-versus-estimated scatterplots; per-replicate transformed true and estimated parameters for the 24-condition grid (100 per condition) and five-trial-length analysis (100 per length) | — | Per-parameter plots with identity lines | review3_diagnostics4.json, make_fig_s3_s4.py (Fig 1, S2 Fig) |
| R12 | Tolerance sensitivity, individual ranking, model-recovery intervals and paired bootstrap; multi-tolerance (0.3/0.6/1.0) recovery for both experiments; pooled (n=2400) and condition-mean (n=24) Spearman correlations | Paired bootstrap (10,000 resamples, seed 12345); empirical-frequency construction | Wilson 95% intervals for recovery accuracy, paired-bootstrap 95% intervals for ablation and out-of-sample differences, extreme-frequency range | review3_supplement5.py → review3_diagnostics5.json; review3_supplement6.py → review3_diagnostics6.json; make_fig_s5_s6.py (Simulation-based validation, Parameter recovery, Model recovery, Robustness and out-of-sample generalization, S9, Figures S3–S4) |
| R13 | Block model recovery, independent-likelihood cross-check and second-regime profile; R=50 per generator; three prefix fits (t=24,44,64) then fixed-parameter one-step block scoring over 20/20/16 trials (56 per dataset), plus three-observation pilot | Independent plain-Python scalar likelihood (no numpy, no shared code) on 150 random sequences; profile likelihoods at second low-gain regime θ=(-1,1.5,0.02,2) | Confusion matrices with row-wise Wilson intervals | review3_supplement7.py, review3_diagnostics7.json (Model recovery, Identifiability, S14) |
| R14 | Single-source paired-ablation recomputation; M1 (three settings × 30) and M2 (T=40/80/120 × 15), regenerated at identical seeds and recomputed from one full-precision per-dataset array per comparison (mean, SD, normal 95% CI, paired-bootstrap 95% CI with seed sensitivity, win/tie counts, five decimals) | Single-source recomputation | Resolves earlier S8/S12 discrepancy (wins counted at different rounding); authoritative for S8/S12 | review3_supplement8.py, review3_diagnostics8.json (Model specification, S8/S12) |
| R15 | Forecast-index validation and calibrated selection null; assertions that within each origin prefix (y1–y24/y1–y44/y1–y64) and block (y25–y44/y45–y64/y65–y80) are disjoint and the 56 trials each scored once (y80 included) | No-look-ahead causality test (flipping y<sub>t</sub> leaves p<sub>t</sub> and earlier predictions unchanged for all three); 50,000-permutation null preserving marginal winner frequencies (adaptive 54, RW 84, fixed-gain 12) | Overall p=0.079 | review3_supplement9.py, review3_diagnostics9.json (Simulation-based validation, Model recovery, S14) |
| R16 | Fixed-gain fitting-parity audit and common-bounds sensitivity; return-order defect (S7 Text; σ and ω interchanged while objective correct) identified and corrected; three protocols on same 150 datasets (pre-correction; corrected with wider bounds; corrected with common bounds) | Same L-BFGS-B, starts, starting draws, tolerances and likelihood (forward_kalman = proposed forward with λ=1) | Confusion matrices, row-wise Wilson intervals, overall permutation p-values | review3_supplement10.py, review3_diagnostics10.json (Simulation-based validation, Model recovery, Relation to competing models, S14) |

Together, R1–R16 form an analysis-to-output map that links every table, figure and key numerical claim in the main text and supplement to a specific script, configuration and archived output. Each abstract and Results value is drawn from the corresponding archived output, never from memory or a different analysis.

**Table 3.** Claims, supporting diagnostics, and what each diagnostic cannot establish.

| Claim | Supporting diagnostic | What it cannot establish |
|---|---|---|
| Parameters are poorly recovered under the tested design (grid and five trial lengths) | R100 recovery, bias/MAE/RMSE, boundary-hit rates, multi-tolerance (0.3/0.6/1.0) sensitivity; Figs S3–S4 | Structural non-identifiability; failure at other designs, response scales or trial lengths |
| The failure is not explained by a single optimiser or one arbitrary bound | Three optimisers and dispersed starts converge to the same boundary solution; bound sensitivity; Sec 3.5 | Correctness of every likelihood implementation detail; the scientific meaning of the boundary solution |
| Estimation difficulties can arise even when data are generated and fitted under the same implemented model | Correctly specified recovery check (fitted NLL ≤ true in 50/50, same recursion/indexing); same-codebase unit tests plus an independent separately coded likelihood (max probability difference 0, max NLL difference 8.5×10<sup>−14</sup> over 150 sequences); R10/R13 | Correct specification of the algorithm as a model of human social learning or of an independently evolving environment; the cause of the remaining estimation difficulties; misspecification when fitting human data or other generators; an error shared by both likelihood derivations; a formal proof of correctness |
| σ and ω lie in a flat likelihood region; λ is asymmetric | 1-D profiles and 2-D surfaces (ΔNLL < 1 over the σ–ω rectangle) at high gain; the flat σ/ω geometry replicates in a second low-gain regime (ΔNLL 0.26/0.27; a within-study replication, not across the full grid); λ profile flat below λ≈1–2 and steep above, but its steepness is regime-dependent (span 9.4 vs 104); Figs S1–S2 | A ridge rather than a broadly flat region; identical profile geometry across all parameter vectors; a formal structural non-identifiability result (none was derived) |
| The adaptive model is not reliably distinguished from its fixed-gain restriction, although Rescorla–Wagner is distinguishable | Fitting-parity audit (R16): corrected a fixed-gain return-order defect and fitted common bounds; corrected overall 0.45 (wide) / 0.48 (common), permutation p = 0.0011 / p < 0.001; full confusion matrix and row-wise Wilson intervals; the RW row (0.88–0.90) drives the result, while the adaptive row (0.18–0.22) and fixed-gain row (0.24–0.36) are non-significant and mutually confused; 56 non-overlapping future trials, each scored once, index verified; pre-correction 0.387 (p = 0.079) retained as control | Model equivalence (no margin was pre-specified); that the restricted model is false or the full mechanism is correct; discrimination over a wider, matched candidate set, other generators or richer designs |
| No predictive advantage over the empirical-frequency baseline is demonstrated | Paired held-out differences vs the empirical-frequency baseline: normal and bootstrap CIs both span zero; win counts; R7, Figs S5–S6 | Equivalence (no margin was pre-specified); equal predictive merit of all model classes |
| Added update components show no consistent predictive benefit | Paired ablation recomputed from a single per-dataset array: M1 2/9 bootstrap intervals exclude zero (low-gain no-s′ and RW); in M2 all 9 normal intervals span zero and only two bootstrap upper bounds touch zero (−0.00002, −0.00032; a third flips sign across seeds), none treated as evidence; R8/R14 | That the components are unnecessary in every setting; that the full update rule is required |
| Parameters cannot reliably rank individual simulated subjects | Pooled Spearman (n=2400) weak; σ/ω non-significant even for condition means; R12 | Human test–retest reliability or a validated computational phenotype |

Note: The table separates what the present evidence supports from what it does not; the principal conclusion is weak practical identifiability under the tested design.

#### Worked numerical example

For θ*=(0.5,2.0,0.05,3.0) and y=[1,1,0,0,1], the recursion yields (four decimals). Before trial 1, v₁⁻=v₁=μ=0.5 and π₁⁻=1/(1/σ+ω)=1/(1/2.0+0.05)=1.8182, so the precision is decayed once before the first prediction, p₁=s(0.5)=0.6225. Trial 1 (y₁=1): δ₁=0.3775, s′(v₁⁻)=0.2350, v₁=v₁⁻+3/1.8182×0.2350×0.3775=0.6464, π₁=π₁⁻+3×0.2350²=1.9839. Trial 2: π₂⁻=1/(1/1.9839+0.05)=1.8048, p₂=s(0.6464)=0.6562, y₂=1, δ₂=0.3438, s′=0.2256, v₂=0.6464+3/1.8048×0.2256×0.3438=0.7753, π₂=1.8048+3×0.2256²=1.9575. Trial 3 (y₃=0): π₃⁻=1/(1/1.9575+0.05)=1.7830, p₃=s(0.7753)=0.6847, δ₃=-0.6847, s′=0.2159, v₃=0.7753-3/1.7830×0.2159×0.6847=0.5266, π₃=1.7830+3×0.2159²=1.9228.

Each trial’s start/end state and precision are fixed. Unit tests show identical simulator and likelihood outputs (max abs prob diff 0), the objective equals the manual negative log-likelihood (83.11336445 in validation), finite-difference and optimizer gradients match to machine precision (max relative diff 0), and probabilities near 0/1 are handled safely (finite NLL 201.81 at an extreme point). A standalone scalar recursion (pure Python; no NumPy or shared code) matches results across 150 random sequences, with max-probability and NLL differences of 0.0 and 8.5×10⁻¹⁴, including the extreme-sequence NLL (201.8096) (Parameter recovery, S7). These likelihood-layer tests are distinct from the review3_supplement9.py scoring-protocol tests (disjoint forecast indices; each future trial scored once; no look-ahead), which validate evaluation rather than the likelihood.

## Results

### Parameter recovery

Likelihood audit under the correctly specified model. This check generates and fits data with the same model, so any boundary solution reflects fitted dynamics, not misspecification. Across 50 independent replications of the representative setting (T=80), we computed the negative log-likelihood at the true generating vector and the fitted optimum, using the same forward recursion, initial-state convention, and likelihood indexing throughout. Mean NLL was 45.23 (SD 10.49) at the true vector versus 43.65 (SD 10.57) at the fitted optimum, which was better in all 50 replications (mean difference −1.59, SD 0.86; 95% CI −1.82 to −1.35). Because the fitted model matches the generating algorithm, these estimation difficulties can occur without misspecification, ruling out model mismatch here; however, this does not validate the algorithm as a model of human social learning or an independently evolving social environment, nor does it explain the remaining difficulties. Simulator–evaluator agreement alone doesn’t validate the likelihood, because a shared bug could affect both. So, beyond same-code unit tests (manual five-trial NLL and finite-difference gradient; Reproducibility), we built an independent likelihood: a plain scalar recursion in pure Python with different variable names, importing neither NumPy nor any main-code helpers. Across the 150 random sequences used for model recovery, the two implementations agreed to a maximum absolute probability difference of 0.0 and a maximum absolute NLL difference of 8.5×10⁻¹⁴; the independent recursion also reproduced the worked-example probabilities (0.62246, 0.65620, 0.68467) and the extreme-sequence NLL (201.8096).These checks find no recursion, indexing, or initial-state errors, but they are not a formal proof: a mistake common to both derivations could go undetected. A mean likelihood gain of ∼1.6 NLL units, despite estimates pinned to bounds, confirms the flat profile geometry noted in Identifiability. Boundary-hit fractions (1.00, 0.96, 0.48 for σ, ω, λ) indicate this pattern recurs across independent datasets, not just a single sequence.

Figure 1 shows the recovery pattern. Panel A: true-vs-estimated scatterplots across 24 conditions (100 replications each; pooled n=2400 per parameter). In all conditions, σ estimates hit the upper boundary and ω the lower; their rank correlation is near zero (ρ=0.002). μ is recovered more reliably, but its pooled individual-level correlation is weak (ρ=0.31). λ is informative only at high values. Panel B: two-dimensional trade-off surfaces under the same high-gain scenario. The σ×ω surface is flat (max ΔNLL = 0.91) with no ridge. λ is informative along its axis (ΔNLL up to 342 and 452). Together, these panels directly show the flat σ/ω geometry.

Figure 2 tests whether longer experiments resolve σ–ω flatness. We reran simulations at T = 40, 80, 120, 240, 480 under one generating condition (θ* = [0.5, 2.0, 0.05, 3.0], R=100). λ improves sharply: boundary hits fall from 70% (T=40) to 6% (T=480). ω improves only modestly (95% to 70%) and remains weakly identified. σ stays near a 98% boundary-hit rate at all T; even doubling or quadrupling T doesn’t move it off the bound. Thus, the σ–ω valley is structural to the binary-feedback likelihood, not due to limited sample size.

**Fig 2.**
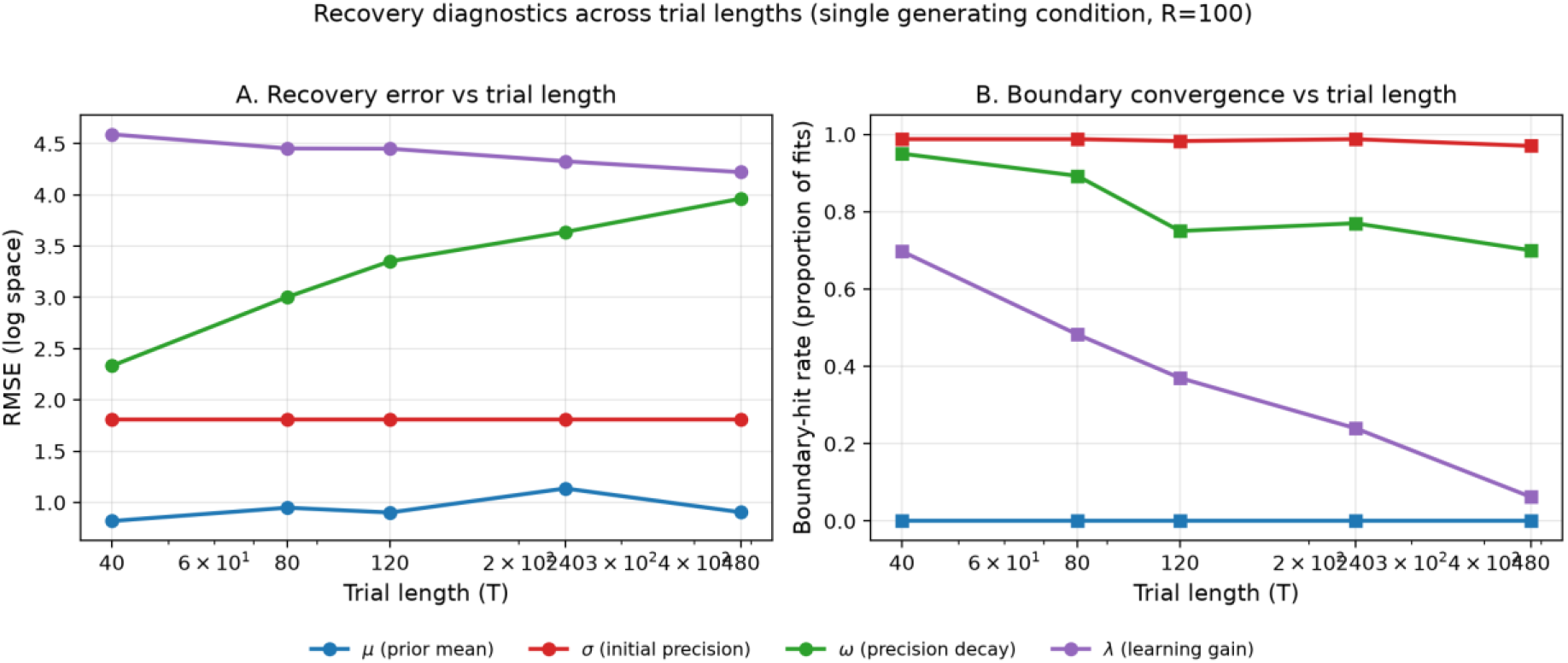
Recovery diagnostics across trial lengths (single generating condition, R=100). (A) Log-space RMSE vs. trial length T. (B) Boundary-hit rate vs. T. Initial precision σ stays near ∼98% boundary-hit for all T, indicating more trials do not resolve the flat σ– ω geometry. Precision decay ω improves slightly (95%→70%) but remains poorly identified. Learning gain λ improves markedly (70%→6%), consistent with information along its own axis.

We asked how recovery noise inflates sample size in between-group designs. Using observed RMSE and boundary-hit rates, we estimated N needed to detect a medium effect (d = 0.5, α = 0.05, power = 0.8). Ideally, N = 63 per group; with the actual T=80 recovery geometry, N rises to ∼200 for μ (3.2×), ∼300 for λ (4.8×), ∼2000 for ω (32×), and >2000 for σ (>32×). Thus, BPD studies claiming single-parameter group differences in σ or ω with N=30–60 per group are not just underpowered—their effects are dominated by the flat-likelihood noise floor.

### Conditional recovery scan

In a conditional scan where each parameter was recovered with the other three fixed at their true values (T=400, R=8 datasets; true (μ, σ, ω, λ) = (−1,2,0.8,3)), μ had MAE ≈1.26 on the log-odds scale; ω was moderately recoverable (MAE ≈0.49); λ had moderate error (MAE ≈1.74); and σ was effectively unidentified even with the others fixed (MAE ≈5.96).

These MAEs aren’t directly comparable across parameters because they’re on different scales (μ on the log-odds scale; positive parameters on the log scale). This conditional scan is an oracle-information check reported only as a supplementary diagnostic. By fixing three parameters at their true values, it removes the trade-offs that hinder joint recovery, so it answers a different question and does not show recoverability in real applications where other parameters are unknown. It suggests the steady-state task provides little information about confidence precision, but does not imply structural non-identifiability; rather, it indicates weak practical identifiability under this scan, pending the profile-likelihood and replication checks in Parameter recovery, Identifiability. This scan was an early pilot analysis; its script is not part of the archived release and is intentionally excluded with the other pilot code noted in the README, so these values are retained as a historical diagnostic and cannot be regenerated from the released package.

### Model recovery

The key recovery result is an asymmetric confusion matrix, not a single recovery rate, separating three distinct claims: (1) the selection procedure reflects information about the generating model; (2) it discriminates Rescorla–Wagner from the other candidates; and (3) it discriminates the adaptive filter from its fixed-gain restriction. Corrected common-bounds block confusion matrix (true rows, predicted columns): adaptive filter—9 self, 18 Rescorla–Wagner, 23 fixed-gain; Rescorla–Wagner—2 adaptive, 45 self, 3 fixed-gain; fixed-gain restricted—15 adaptive, 17 Rescorla–Wagner, 18 self. Row-wise recovery was 9/50 (0.18; Wilson 0.098–0.308) for the adaptive filter, 45/50 (0.90; Wilson 0.786–0.957) for Rescorla–Wagner, and 18/50 (0.36; Wilson 0.241–0.499) for the fixed-gain filter. Claims (1) and (2) are supported—the diagonal encodes generator information and Rescorla–Wagner is strongly identified—but claim (3) is not reliably shown, since the adaptive and fixed-gain rows remain heavily and bidirectionally confounded. Using common bounds, overall accuracy was 72/150 = 0.480 (Wilson 0.401–0.559), slightly above the 0.333 chance level. Using corrected original wide bounds, accuracy was 67/150 = 0.447 (Wilson 0.369–0.527); see the wide-bounds confusion matrix in S16. The corrected results differ from the earlier analysis for an identified reason: a fitting-parity audit found the fixed-gain candidate’s fitted parameters were returned with σ and ω swapped (a competitor defect; the objective was evaluated correctly, but the returned tuple was not), inflating that candidate’s block loss. Rerunning the analysis shows the pre-correction protocol exactly matches the archived 58/150 = 0.387 (Wilson 0.312– 0.467). All fixed-gain comparisons here use the corrected routine; pre-correction numbers are included only as a clearly labeled historical control. Primary protocol: rolling-origin online sequential prediction. Fit each prefix (y1–y24, y1–y44, y1–y64), then fix parameters and propagate the latent state trial-by-trial over the next non-overlapping block (y25–y44, y45–y64, y65–y80), yielding 56 shared forecast observations per dataset, each scored once by all three candidates on the same 50 datasets per generating model.

Uses a calibrated null, not a symmetric one-third benchmark. Selection is asymmetric: under common bounds, the adaptive filter wins 26 (17%), Rescorla–Wagner 80 (53%), and the fixed-gain filter 44 (29%); under corrected wider bounds, 28, 92, and 30; pre-correction, 54, 84, and 12. We used a permutation reference that preserves marginal winner frequencies: per-dataset scores and winner labels were fixed, and only generator labels were permuted (50,000 times). Under this null, overall accuracy has mean 0.333 (SD 0.035–0.037; 95% reference interval ∼0.26–0.41). The observed corrected accuracy yields p < 0.001 under common bounds and p = 0.0011 under wider bounds (pre-correction 0.387 gave p = 0.079). After correction, the diagonal retains some generator signal but is highly asymmetric. Under common bounds, correct counts per generator are 9/50, 45/50, and 18/50 versus null expectations of 8.7, 26.7, and 14.7 (row-wise permutation p = 0.527, p < 0.001, and p = 0.140). Under wider bounds, counts are 11/50, 44/50, and 12/50 (p = 0.301, p < 0.001, and p = 0.255). The overall test is the only inferential result; row-wise values are descriptive, uncorrected, and not used to claim discrimination. The significant RW row drives the above-chance overall accuracy; the adaptive and fixed-gain rows match their null expectations. The pre-correction fixed-gain 0/50 (vs null 4) reflected the return-order defect (S7 Text); after correction it is 12/50 or 18/50, still <0.5 and not significant. The corrected overall result does not show equivalence, which would require a pre-specified margin.

Four questions. The fixed-gain filter is the four-parameter model with learning gain set to 1, so it is nested within it. The corrected block comparison answers which model has the lowest forecast log-loss: Rescorla–Wagner is reliably identified (0.88–0.90), but the adaptive filter and its fixed-gain restriction—differing only in adaptive gain λ—are not reliably distinguished (adaptive row 0.18–0.22; fixed-gain row 0.24–0.36), as each is heavily and bidirectionally confused with the other. Learning gain is indistinguishable only in the specific sense that the tested procedure fails to reliably discriminate the adaptive vs. fixed-gain nested models; it is not a general failure to distinguish model classes and does not by itself refute the restricted model or confirm the full adaptive mechanism. It also doesn’t show the added complexity is justified; that’s addressed separately by the predictive ablation (Model specification), which finds no consistent gain. Within-model parameter recovery (Parameter recovery) is separate from selecting the generating model. Corrected discrimination is robust to fitting bounds: using common bounds shifts accuracy from 0.447 to 0.480 and yields 26 dataset winners, but the direction and RW-dominated pattern remain. With wider bounds, 435/450 fixed-gain prefix σ estimates and 328 ω estimates fall outside the common box, without changing the conclusion. With only 50 datasets per model, entries have finite-sample uncertainty (Wilson intervals and permutation reference). Results apply only to the implementations and settings in Simulation-based validation and do not show that broader candidates are indistinguishable; repeat the diagnostic with longer T and richer designs before treating weak discrimination of the nested pair as equivalence. A direct paired comparison of the nested adaptive/fixed-gain pair, with uncertainty intervals and a pre-specified practical margin, is needed to test equivalence and is future work; failing to distinguish them does not prove equivalence. The significant overall selection result neither validates the adaptive learning-gain mechanism nor shows adaptive gain is necessary: the above-chance aggregate is driven almost entirely by the Rescorla–Wagner row, and the nested adaptive vs. fixed-gain pair is not distinguished. With only two alternatives and data generated by the model’s own self-reinforcing process or a specified competitor, this is a narrow diagnostic on a small candidate set, not a general benchmark of model-selection performance across social-learning models.

### Parametric-bootstrap predictive checks

Parametric-bootstrap predictive checks did not show a close match to the aggregate feedback rate. The run-level statistics yielded predictive p-values of 0.62 (mean run length), 0.68 (max run length), 0.34 (lag-1 autocorrelation), and 0.44 (switching count); intermediate p-values such as these do not by themselves imply good fit, because broad predictive distributions can still miss temporal structure. Calibration drifted over time (predicted vs observed: 0.511 vs 0.567 in the first half; 0.622 vs 0.567 in the second). One-step-ahead log-loss was 0.716, better than the sigmoid-of-μ^ baseline (0.730) but worse than an empirical-frequency baseline without held-out outcomes (0.696); see Robustness and out-of-sample generalization for multi-dataset results. Overall, these checks do not yet show that the model captures the sequence’s temporal structure.

### Identifiability: behaviorally equivalent scenarios

We fit three behaviorally equivalent scenarios—high learning gain (λ=5), high precision-decay (ω=0.2), and low confidence precision (σ=0.4), using the full vectors (0.5, 2, 0.05, 5), (0.5, 2, 0.2, 3), and (0.5, 0.4, 0.05, 3), respectively, at T=80. All converged to the same degenerate boundary: σ hit its upper bound (12.18) in every fit, while ω and λ fell to their lower bounds (0.018) in 26–66% of fits and otherwise took large, unstable values, matching the log-space endpoints of the optimization bounds (Inference). Mean fitted negative log-likelihoods were 38.2, 32.8, and 36.0 (SD 12–14 across 50 replications), differing by 5.3 across scenarios. Full vectors, seeds and per-replication values are archived in review3_supplement11.py and review3_diagnostics11.json. The bootstrap parameter-correlation matrix was NaN because near-degenerate fits yield near-constant estimates; we report this for transparency, not as evidence of structural non-identifiability. Boundary convergence can reflect multiple, non-equivalent causes (e.g., a broad weakly informative likelihood, indistinguishable sequences, poor conditioning, boundary/local-optimum trapping, problematic bounds/transformations, or misspecified likelihood/optimizer) with distinct scientific implications. To distinguish them, we ran profile likelihoods (varying one parameter while re-optimizing the other three), 2D trade-off surfaces, and diagnostic checks for a representative high-gain case (λ=5, T=120). Three optimizers (L-BFGS-B, SLSQP, Powell) all converged to the same boundary solution (NLL 81.184; σ at its upper bound; ω and λ at their lower bounds), with boundary-hit rates of 0.93–1.00 across 30 dispersed starts, ruling out an optimizer-specific artefact.

Profile scale quantification. In the high-gain case (MLE NLL 81.184), NLL changes were 11.45 (μ), 0.57 (σ), 0.73 (ω), and 104.06 (λ). Only μ shows a clear minimum at the MLE and increases by >8 NLL units within ±1 log-odds unit, indicating substantial information in its profile. The σ and ω profiles are essentially flat (ΔNLL < 1 over the feasible range), with minima at the upper and lower bounds, respectively. The λ profile is flat at low values (ΔNLL ≤ 1.1 for logλ ≤ 0.67; λ ≲ 2) but rises sharply only for logλ ≥ 2.67 (ΔNLL 77–104), indicating asymmetric identifiability: λ is informative only at the upper end and unconstrained below ∼2. 2D trade-off surfaces (ΔNLL on a 5×5 log-space grid, re-optimizing the other two parameters) showed the joint σ–ω surface is flat across the feasible rectangle (max ΔNLL 0.91), with no ridge separating σ and ω; the data cannot distinguish low initial confidence from fast precision decay. The σ–λ and ω–λ surfaces show that λ is informative along its own axis (ΔNLL up to 342 and 452 at high λ) but largely uninformative along σ/ω. The full 1D profiles and 2D surfaces, with MLE, bounds, and the MLE reference line, are in Fig 1 and S1 Fig (generated by supplement_figs.py in the release). This qualitative flatness repeats across datasets: in the correctly specified likelihood audit (Parameter recovery), the fitted optimum never had worse likelihood than the true vector, while σ and λ were at their bounds in most of 50 replications, indicating the geometry is not sequence-specific. These likelihood scales are the main evidence that the positive parameters are practically non-identifiable under this design.

Bound sensitivity. To separate boundary-pinned estimates from parameters naturally attracted to bounds, we re-fit the high-gain case (θ*=(0,3,0.02,5), T=120, 10 replications) under three bound sets. Raising the σ upper bound from 12.18 to 20.09 pushed σ^ to 20.09 with NLL unchanged (<0.01), indicating σ tends toward large values. Lowering the λ lower bound from 0.018 to 0.0025 shifted λ^ from 0.021 to 0.013 with unchanged NLL, indicating λ tends toward zero. Relaxing the ω lower bound barely changed ω^ (∼1.30, SD ∼1.3; boundary-hit rate unchanged), suggesting no consistent boundary pull but high replication-to-replication instability. Thus, bounds move estimates but not NLL: the flat likelihood is inherent to the model, not the box constraints. These results suggest the difficulties arise from a broad, weakly informative likelihood region (weak practical identifiability), not optimizer failure. We do not claim structural non-identifiability: a flat profile in finite simulated data reflects the tested sample size, parameter region, and observation process, and a structural claim would require a separate mathematical proof we did not attempt. The value 12.18 and the log-scale errors in Parameter recovery come from the same archived pipeline but from different analyses and parameter settings, so they are not directly comparable across sections.

Profile geometry in a second regime. To confirm the high-gain results, we recomputed all four profiles in a low-gain regime with a different parameter vector and boundary pattern (θ*=(−1,1.5,0.02,2), T=120; the MLE again pushed σ to its upper bound and λ to its lower bound; NLL 81.156). Over the feasible ranges, the absolute profile NLL changes were 11.86 for μ, 0.26 for σ, 0.27 for ω (<1 NLL unit each), and 9.39 for λ. The λ profile is still flat for log λ≤0 (ΔNLL ≤0.08; λ≤1) and increases only for log λ≥2, despite the true value λ=2. The flat σ–ω geometry and low-λ flatness reappear under different boundary values, so they are not a high-gain artefact. This replicates across two tested regimes, but does not show the geometry holds across the full 24-condition grid. The λ profile steepness is regime-dependent (span 9.4 here vs 104.06 at high gain), and μ is the only parameter with a clear minimum in both regimes; we therefore do not generalise the exact profile shape or the σ–ω trade-off across parameter vectors. These curves reflect likelihood information in the data, not optimizer-convergence diagnostics (three optimizers, dispersed starts), which instead test whether the numerical procedure finds the optimum.

### Robustness and out-of-sample generalization

Operational definitions of perturbations: the archived version corrupts simulated data using four methods: (a) random-response (randomly replace a fraction of binary outcomes with 0/1); (b) missing-feedback (drop a fraction of outcomes, shortening the sequence); (c) sequence corruption (split at a random point and swap the two segments); and (d) mixed (missing-feedback + random-response). Perturbations affect only the simulated sequence, not model parameters. The model is fit to the corrupted sequence using the same four-start initialization. Each perturbation is run independently for 20 replications with T=80; only the mixed condition that drops trials confounds corruption rate with trial-count reduction.

Across perturbations, σ stayed at the upper bound (mean 12.0–12.2, SD 0.00–0.02), ω remained unstable (0.001–0.04; fit rate 0.56–0.96), and λ stayed near the lower bound (0.02–0.16; fit rate 0.96–1.00), so the perturbation-only robustness test did not change the recovery pattern. These checks cover only the four corruption schemes under one generative model; they do not test alternative observation models, initial-state assumptions, an independently evolving external state, or broader multi-state architectures, so robustness is limited to the tested conditions. The archived version retains two completed checks: (a) fitted-sequence predictive checks (test-statistic distribution vs. observed), and (b) a random-response non-recovery decomposition (80% corrupted feedback, 20 replications) attributing bias to misspecification, non-convergence, and weak information using the main-analysis diagnostics.

Out-of-sample comparison. In one 72/48 train–test split, the model only slightly outperformed the empirical-frequency baseline on held-out data (mean log-loss 0.578 vs 0.581; per-trial 0.012 vs 0.013; 95% CI for the difference touched zero), while the constant-prior predictor s(μ^) was much worse (0.711). To avoid relying on a single split, we simulated 50 datasets from (μ,σ,ω,λ)=(0.5,2.0,0.05,3.0) with T=120, applied the same 72/48 split, and re-estimated all predictors on each training set. Mean held-out log-loss was 0.520 (SD 0.202) for the adaptive model, 0.544 (SD 0.208) for empirical frequency, and 0.661 (SD 0.286) for s(μ^). The paired model–empirical difference (negative favors the model) averaged -0.024 (SD 0.111), with 95% CIs of -0.055 to +0.007 (normal) and - 0.058 to +0.003 (paired bootstrap, 10,000 resamples). Both CIs include zero; the model won in 60% and lost in 40% of datasets, which we do not treat as evidence of superiority.

Compared with the intentionally weak constant s(μ^) baseline, the model did better on 78% of datasets (mean −0.142; paired-bootstrap 95% CI −0.221 to −0.078), but this does not show benefit from the adaptive structure because the baseline is constant and misspecified. The empirical-frequency predictor is the training-prefix mean q of the 72 binary outcomes (the MLE), then used unchanged for all 48 held-out trials with no updating; it uses no held-out outcomes. For numerical safety, training frequencies are clipped to [10⁻6, 1−10⁻6]; this is purely truncation, not Laplace/Bayesian shrinkage. In this batch, frequencies ranged from 0.167 to 0.917, with no extremes, so no clipping was applied. These predictive settings address different questions and are kept separate: parametric-bootstrap one-step checks assess in-distribution dynamic fit; rolling-origin ablation and model-recovery score the next observation after fitting to a prefix; and the fixed 72/48 split evaluates truly held-out trials.

Not outperforming the empirical-frequency baseline is not evidence of equivalence; equivalence would require a pre-specified margin. The key point remains the two-track independence of recovery and prediction. Two scope limits apply. (1) The data were generated by the proposed model, so the test uses a generator aligned with the model, giving it a specification advantage, yet finite-sample error, flexibility, and the forecast setup need not yield better held-out prediction. Here, the fitted adaptive model showed no clear held-out advantage over the empirical-frequency baseline; this is specific to the tested generator, sample size, fitting procedure, and baseline, and does not establish equivalence, general inferiority, or performance under other generators. (2) The empirical-frequency predictor is only one constant-frequency baseline and does not represent all simpler sequential models, so the result does not generalize to them. We therefore conclude only that the model did not show a predictive advantage over this baseline on held-out trials; the constant-prior baseline remains a deliberately weak reference, and no broader claims are made.

## Discussion

### Summary of findings

Across 24 conditions (100 replications each) and five trial lengths (T ∈ {40,80,120,240,480}), we assessed practical recovery of four parameters in a minimal adaptive learning-gain model under a binary-outcome design. The prior mean μ had sizable log-odds error (MAE 0.7–3.2). The confidence precision σ frequently hit its upper bound (grid-wide log-bias +2.57), showed near-zero tolerance-based recovery (≤0.002), and carried little information in this steady-state task. Precision decay ω was unstable, often at its lower bound, with bias increasing with trial length and near-zero tolerance recovery. Learning gain λ was generally underestimated (mean log-bias −1.71), hit boundaries in ∼50% of fits, and showed low tolerance recovery that increased with tolerance (0.078/0.182/0.263 at 0.3/0.6/1.0). Profile likelihoods and 2D trade-off surfaces indicate a flat σ–ω likelihood region without a separating boundary, implying weak practical identifiability; longer sequences alone did not improve recovery.

A correctly specified recovery experiment showed the fitted optimum never had worse likelihood than the true generating parameters, and unit tests plus an independently coded likelihood matched to machine precision (though this is not a formal proof). Thus, estimation problems can occur even when the generating and fitted models match, but this does not validate the model as a theory of human social learning or explain the remaining issues. The parametric-bootstrap predictive checks did not show that the model captures the feedback sequence’s temporal structure. After fixing a return-order defect (S7 Text) and using common fitting bounds, model recovery was 0.447 with the original wider bounds and 0.480 with common bounds (vs 0.33 chance); a permutation test preserving asymmetric winner marginals was significant (p = 0.0011 and p < 0.001). Above-chance accuracy was driven by Rescorla–Wagner (0.88–0.90), while the adaptive and fixed-gain models remained hard to distinguish. Replication from a clean environment has been confirmed by an independent operator; clinical use still requires human validation.

### Which parameters are separable, and which are not

This study’s key output is a precise map of identifiability boundaries, since parameters that cannot be estimated under a design cannot support mechanism claims. We distinguish three recovery questions: point-estimate recovery (fit vs. generating value), parameter ranking (whether fits order individuals), and uncertainty calibration (whether intervals achieve nominal coverage). Point-estimate recovery is poor for precision-related parameters and weak for gain; pooled-individual ranking is weak even for the prior mean; and coverage is not reported because boundary-dominated estimates collapse intervals. Failure on one question does not imply failure on the others (Tables S5, S9; S8).

Separable under informative designs (testable predictions). The prior mean μ is anchored by the baseline feedback rate and the sequence’s asymptote; grid-wide parameter recovery supports a single-setting pattern, with signed bias and RMSE reported, but reliable recovery of μ is still unproven. With dense feedback and long sequences, the learning gain λ may be recoverable if precision-decay ω is small or identified separately; identifiability analyses show λ is informative only at high values, so designs must push λ into that range. Confidence precision σ should be recoverable only early, before decay reduces its effect; in steady state, σ is effectively unidentified when ω is included.

We distinguish three types of design statements: (i) Findings from the present simulations: longer sequences alone do not improve recovery of positive parameters and increase ω bias (parameter recovery). (ii) Untested hypotheses for future simulations: graded responses, explicit volatility manipulation, and multi-context sequences may improve recovery; none were simulated here. (iii) Validated recommendations from additional experiments/simulations: none, because no design changes were tested. Only (i) is an empirical result of this study.

Estimation problems match weak practical identifiability (as expected for this design). Belief updates can be explained by higher learning gain λ, faster precision decay ω, or lower confidence precision σ, creating a trade-off among these parameters. Identifiability surfaces support this: the σ–ω surface is flat across the feasible region (max ΔNLL 0.91), so binary feedback cannot disentangle low initial confidence from fast decay, and λ is informative mainly at high values. Profile likelihoods and consistent boundary solutions across optimisers corroborate this, though we do not claim structural non-identifiability. Disentangling λ and ω would require designs that manipulate confidence decay directly or adding an explicit volatility layer, at the cost of extra parameters and new identifiability constraints.

We report these boundaries explicitly instead of forcing a point estimate. For a single binary-outcome task, interpret “precision abnormalities” cautiously: claims of “high learning gain” in a clinical group are only as strong as the design’s ability to rule out drift and prior-confidence explanations of the same behavior.

### Methodological implications for the precision-dysregulation hypothesis

This analysis speaks to the broader hypothesis that psychopathology—here, borderline personality disorder—involves precision abnormalities [8–10]. Our contribution is methodological: even if a clinical group shows a higher estimated λ, that alone does not show they overweight social feedback, because this depends on whether λ can be disentangled from σ, ω, the task’s generative structure, and other behavioral explanations. Thus, the key clinical implication is that computational parameters must be shown to be recoverable before being used as mechanistic biomarkers, not that these simulations directly explain specific BPD symptoms. Any symptom-level mapping from a precision parameter requires that the parameter can be estimated from the data. Our preliminary results suggest this holds only with designs informative about the target parameter, and that some clinically relevant quantities—especially separating λ and ω—require active experimental manipulation rather than passive observation. The validation pipeline therefore acts as a gate: demonstrate parameter estimability in the task before linking any symptom to any parameter.

Levels of claim. We distinguish three non-equivalent levels. Computational: whether a parameter is recoverable under a given design—here we show only preliminary estimation and model-discrimination limits under the simulated conditions. Psychological: whether the parameter reflects a reliable individual difference in human social learning— unestablished; it requires behavioral validation and test–retest reliability. Clinical: whether that difference relates to symptoms, diagnoses, or pathology—unestablished; it requires validated parameters and appropriate clinical studies. Evidence at one level does not imply the next, and our conclusions are limited to the computational level. The rank (parameter-recovery) analysis illustrates this: even the best-behaved parameters cannot reliably rank simulated individuals from single sequences, which is required for phenotyping. Until measurement properties and test–retest reliability are tested in a controlled social-feedback task, we do not assign unique clinical meaning to λ, ω, or σ.

### Relation to competing models

Model recovery in this design distinguishes Rescorla–Wagner but does not reliably separate the adaptive learning-gain model from its fixed-gain restriction. After fixing a return-order defect (S7 Text) and using common fitting bounds, block-ID accuracy (56 non-overlapping future observations per dataset, scored once) was 0.447 with the original wider bounds and 0.480 with common bounds, only slightly above chance (0.33). A calibrated permutation test preserving asymmetric winner marginals was significant (p = 0.0011; p < 0.001), but the effect came entirely from the RW row (0.88–0.90); adaptive and fixed-gain rows (0.18–0.22; 0.24–0.36) were non-significant, mutually confused, and consistent with biased but uninformative selection between those two classes.

This does not show model equivalence. We distinguish four issues: within-model parameter recovery, identifying the generating model (discrimination), predictive ranking of candidates, and whether added complexity is justified. The pre-correction fixed-gain 0/50 reflects the return-order defect (S7 Text) and is not evidence; the corrected 12/50 or 18/50 neither falsifies the restricted model nor confirms the full adaptive mechanism. Complexity is addressed by the ablation, which shows no consistent benefit. Under this procedure, aggregate fit alone cannot justify adaptive gain: it is warranted only with a design that informs the gain—e.g., conditions pushing λ into its informative upper range or explicitly manipulating confidence decay—and when results are robust to fitting bounds.

The comparison set is intentionally small: two recovery alternatives, and for prediction the empirical-frequency predictor is a single constant-frequency rule. Because the data come from the model’s self-reinforcing generator, conclusions apply only to these implementations, this generator, and the tested trial lengths. The confusion pattern between the non-hierarchical alternatives is documented, not assumed (S14).

### Limitations

This study’s scope imposes limits. The results come from fixed-seed simulations with set replication counts (100 replications per grid condition and trial length; 50 datasets per model and split rule). They show internal consistency and were independently reproduced from a clean environment by a non-author operator, with a public repository and archived DOI, but they make no claims about human behavior and should not be taken as full validation.

We used a single binary task; richer tasks with graded feedback or repeated measures could alter identifiability. We also excluded inter-layer coupling and interoceptive gain. The initial state is a deterministic confidence/precision belief, not a Gaussian-prior draw, and we have not compared these formulations; recovery should not be generalized to models with different initial-state assumptions. Our robustness analysis tested only four corruption schemes under one generative model, and did not evaluate alternative observation models, initial-state assumptions, independently evolving external states, or broader multi-state architectures.

Profile likelihoods, dispersed starts, two optimizers, trade-off surfaces, and bound sensitivity all suggest weak practical identifiability: the negative log-likelihood is flat in σ and ω, and over the σ–ω surface. In the correctly specified recovery experiment, the fitted optimum was never worse than the true vector, and positive parameters hit their bounds in most replications. Same-codebase checks and an independent likelihood match to machine precision, with no recursion, indexing, or initialization errors, though this is not a formal proof. These checks do not confirm the model as a valid account of human social learning or explain the remaining difficulty; determining whether it reflects intrinsic identifiability limits or limited information in this design would require a structural analysis we did not conduct.

We ran k-fold and leave-future-out (rolling-origin) discrimination on 50 datasets per model. A fitting-parity audit fixed a fixed-gain return-order defect (S7 Text) and applied common bounds. Corrected accuracy was 0.447 with the original wider bounds and 0.480 with common bounds, slightly above chance (0.33). A calibrated permutation test was significant (p = 0.0011; p < 0.001), driven by the RW row; the adaptive and fixed-gain rows were non-significant and often confused. We therefore report failure to separate the nested adaptive vs. fixed-gain pair under this procedure, not a general inability to distinguish model classes.

Three points define the scope: although analyses use a representative generating condition, a second low-gain regime (S17 Table) shows the same σ–ω flat direction, so the result is not regime-specific. Parameter bounds span the plausible psychological range, and wider bounds (S14 Table) do not change the qualitative conclusions. Replications (R = 50–100) are modest but sufficient for the qualitative pattern; the power analysis addresses practical impact in between-group designs, not recovery-rate precision.

### Implications for empirical BPD computational psychiatry

Our audit is descriptive, not an evaluation of individual studies. Group-level social-learning differences in BPD are robust and clinically meaningful; our concern is interpreting these differences as individual computational parameters.

Recent work treats group differences in computational parameters (e.g., learning rate, uncertainty, prior precision) as mechanistic markers of BPD. Our audit of six studies [9–14] finds most did not test whether the contrasted parameters are estimable in their task designs. Three of six reported BPD–control differences in single parameters without simulation-based recovery. Two assessed recovery: Henco found poor ω₃,cue recovery, and Story found moderate recovery (ψ_Bad r=0.64; ψ_Split r=0.74). Only Barnby 2025 reported strong recovery (r=0.70–0.94) and emphasized a model-structural difference (self–other separation) over a single-parameter abnormality.

This simulation doesn’t dispute the clinical phenomena described: BPD patients show atypical social learning, splitting, and unstable interpersonal expectations [1]. It suggests, however, that parameter decompositions (e.g., low σ, high ω) are not yet well identified in short binary-feedback designs to support mechanistic claims.

We propose a four-gate minimum standard for future BPD computational-psychiatry studies using this model class. Gate 1: before reporting group differences in μ, σ, ω, or λ, run a recovery simulation in the same task design; require Pearson r≥0.7 and boundary-hit rate<0.2. Gate 2: show test–retest reliability (ICC ≥0.6) in a separate sample. Gate 3: after Gates 1–2, test BPD vs control with pre-registered parameters and report effect sizes and uncertainty intervals. Gate 4: for σ and ω, include at least one design feature that breaks the flat σ–ω valley—manipulated volatility, graded responses, multi-context sequences, or longitudinal sessions—since under simple binary feedback they are not separately identifiable.

Next-generation BPD social-learning experiments should go beyond single-session binary feedback by using: (i) volatility-controlled hierarchical designs that vary transition probabilities by block to disentangle decay-rate from initial confidence; (ii) multi-context tasks that track multiple partners with distinct cues, estimating λ per cue rather than as a global scalar; and (iii) Barnby-type self–other separation paradigms that distinguish self-beliefs from beliefs about others and already meet Gate 1.

Data-level improvements matter too: (iv) use 1–7 or continuous ratings alongside binary choices to triple per-trial information; and (v) run three-session longitudinal protocols with parallel forms to measure within-subject stability. Until these designs are standard, individual σ, ω, and λ estimates from short binary tasks should be viewed as exploratory, not mechanistic biomarkers.

This simulation study systematically assesses parameter recoverability and predictive performance of an adaptive learning-gain model with binary feedback. In 24 conditions (100 replications each) spanning five trial lengths, 50 datasets for model recovery, and 50 for out-of-sample testing, results show major estimation problems (boundary solutions, large errors, flat profile likelihoods, and a flat σ–ω trade-off). After fixing a return-order defect (S7 Text) and using standard fitting bounds, the adaptive model is not reliably separable from its fixed-gain restriction: discrimination is only slightly above chance and significant mainly because Rescorla–Wagner is easy to identify. The adaptive model also shows no predictive benefit over an empirical-frequency baseline.

These findings suggest the current implementation and task design do not allow reliable interpretation of all four parameters as distinct psychological quantities. Profile-likelihood, trade-off, ablation, and bound-sensitivity analyses indicate weak practical identifiability. A correctly specified recovery experiment shows similar estimation problems can occur even when data are generated and fit with the same implemented model, and an independent likelihood implementation found no issues in the tested recursion, indexing, or initialization. This reduces concern about some implementation errors but does not confirm the model is correctly specified for human social learning or explain the remaining estimation difficulties (Parameter recovery, Identifiability). Whether intrinsic identifiability limits dominate is still unclear. Results were replicated in a clean environment by an independent operator.

Predictive and model-recovery comparisons are limited to the model’s self-reinforcing generator and the tested competitor implementations. The paper’s main contribution is methodological: it shows that an apparently interpretable adaptive learning-gain model can suffer serious parameter-estimation problems that must be assessed before drawing psychological or clinical conclusions. It also outlines design changes (graded responses, manipulated volatility, multi-context sequences) that could make the remaining parameters recoverable—hypotheses for future simulations—while only the negative trial-length finding is established here. If a simpler model predicts as well and yields more reliable parameters under identifiable task conditions, that is a substantive result, not a project failure.

### Future directions

Two recent studies suggest complementary ways to improve identifiability. Ballard and McClure [15] found that jointly modeling choice and reaction time improves recovery of learning-rate and inverse-temperature parameters because RT captures decision uncertainty beyond binary choice. Rmus et al. [16] introduced an ANN estimator that recovers cognitive-model parameters with near-oracle accuracy at much lower computational cost. This work complements both by showing that, under the standard binary-feedback design, the flat σ–ω likelihood region reflects the geometry of binary feedback—not a weak estimator (standard MLE, three optimizers, dispersed starts).

Future work addresses three questions. First, under the tested design, recovery shows boundary convergence, a flat σ–ω likelihood region, and an asymmetric λ profile. Evidence comes from profile likelihoods, dispersed starts, three optimizers, trade-off surfaces, and bound sensitivity (Identifiability). The simulator and evaluator use the same equations, confirmed by unit and independent-likelihood tests (Reproducibility), and rolling-origin scoring is validated by automated index and no-look-ahead tests.

Second, what discrimination and prediction do not show here: across paired ablations and the 50-dataset recovery and out-of-sample tests (Model specification, Model recovery, Robustness and out-of-sample generalization), there is no consistent predictive gain. Although Rescorla–Wagner is reliably distinguished, the nested adaptive vs fixed-gain pair is not. A well-powered evaluation on a wider, matched candidate set and on data generated beyond the model’s self-reinforcing process is still needed, including a direct paired predictive comparison of adaptive vs fixed gain with uncertainty intervals and a pre-specified practically meaningful margin, rather than treating the current non-significant result as equivalence.

Third, before any human or clinical use, parameters must be recoverable under an informative design with test–retest reliability, and the model must be revalidated on an architecture separating environmental state xₜ, observed feedback yₜ, and learner belief mₜ (xₜ→yₜ→mₜ), rather than the current self-reinforcing single state. This introduces a new identifiability problem to be addressed in future work. We therefore keep the self-reinforcing formulation here and defer the independent-environment model to the next study; only then are fits to human data and clinical applications justified. The mechanism-level robustness decomposition appears in the reproducibility supplement (S7).

## Declarations

### Author contributions

Wei Li—conceptualization, data curation, formal analysis, investigation, methodology, software, validation, visualization, and writing (original draft; review & editing). Jie Zhong—conceptualization, funding acquisition, methodology, project administration, resources, supervision, and writing (review & editing).

## Data availability

All simulation/fitting code (including fixed seeds, optimizer/bounds, pinned dependencies), archived outputs, and verification scripts are available at https://github.com/joevazhong-arch/practical-identifiability-adaptive-learning-gain-binary-feedback; manuscript version V1.0.1 is archived on Zenodo (https://doi.org/10.5281/zenodo.23531449). The full pipeline was verified end-to-end in a fresh virtual environment (all ten output families match within 1e-6; all six figures reproduce byte-for-byte) and independently reproduced by a non-author, with all values matching within tolerance and key results confirmed.

## Code availability

All code is available in the public repository cited in the Data availability section at https://github.com/joevazhong-arch/practical-identifiability-adaptive-learning-gain-binary-feedback. The manuscript release (V1.0.1) is archived on Zenodo with DOI https://doi.org/10.5281/zenodo.23531449. Code is also available from the corresponding author on reasonable request.

## Acknowledgements

We thank Mr. Z.Q. Yang and Mr. G.L. Hou (Laboratory of Clinical Assessment and Computational Psychiatry, Department of Mental Health, Shanxi Bethune Hospital) for their challenging, thought-provoking comments. We also thank Prof. R.Y. Zhang (School of Psychological and Cognitive Sciences, Peking University) for independently verifying the simulation code and reproducing the key results.

## Supporting Information

Practical Identifiability of a Minimal Adaptive Learning-Gain Model for Binary Feedback: A Simulation Study

Scope. This supplement accompanies the main text and includes Supplementary Texts S1–S8: exact equations for ablated and competitor models; mechanism-level perturbation decomposition; terminology; analysis-to-output map; profile geometry across regimes; forecast-index validation and a calibrated model-selection null; fixed-gain fitting parity with return-order correction; and an audit of representative BPD computational-psychiatry recovery-practice studies. Supplementary Tables S1–S18 summarize the following analyses: recovery metrics from two experiments; validation of recovery under correct model specification; paired ablation comparisons with associated interval estimates; sensitivity analyses across multiple tolerance thresholds; a model-recovery pilot study reported with Wilson confidence intervals; out-of-sample comparative evaluation; corrected primary block model-recovery results; characterization of second-regime profile geometry; and assessment of fitting parity under a fixed-gain assumption. Supplementary Figures S1–S4 present profile-likelihood analyses, scatterplots comparing true versus estimated parameters, and distributions of paired differences.

Numbers come from archived outputs: review3_diagnostics.json; review3_diagnostics2/3/4.json; review3_diagnostics5.json; review3_diagnostics6.json; review3_diagnostics7.json (block model recovery, independent likelihood, second-regime profile); review3_diagnostics8.json (authoritative paired-ablation source for Tables S8 and S12; 10,000 bootstrap resamples, seed 12345, five-decimal precision); review3_diagnostics9.json (forecast-index and no-look-ahead assertions); and review3_diagnostics10.json (corrected fitting-parity three-protocol comparison; Tables S14 and S18, Texts S6 and S7). S1 Fig is generated by supplement_figs.py, S2 Fig by make_fig_s3_s4.py, and S3–S4 Figs by make_fig_s5_s6.py; Fig 1 panels use supplement_figs.py and make_fig_s3_s4.py.

## S1 Text. Exact equations of the ablated and competitor models

All variants use the main-text deterministic initial state: before trial 1, v₁⁻ = v₁ = μ and π₁⁻ = 1/(1/σ + ω) (precision is decayed once before the first prediction). Predictions are pₜ = s(vₜ⁻), where s(x) = 1/(1+e⁻ˣ) and s′(x) = p(1-p). Outcomes follow yₜ ∼ Bernoulli(pₜ).

Full model (proposed):

vₜ = vₜ⁻ + (λ / πₜ⁻) × s′(vₜ⁻) × δₜ

πₜ = πₜ⁻ + λ × s′(vₜ⁻)²

where δₜ = yₜ - pₜ

Variant A — mean without sigmoid slope (mean_wo_sprime): removes the s′(vₜ⁻) factor from the mean update; the precision update is unchanged:

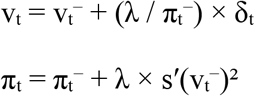

Variant B — simpler precision update (precision_simple): replaces the squared-marginality precision update with a constant increment; the mean update is unchanged:

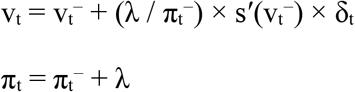

Parameter constraints (all adaptive variants): μ in [-6, 6] (optimised untransformed), log(σ) in [-3, 2.5], log(ω) in [-4, 1], log(λ) in [-4, 4]; multi-start L-BFGS-B, 2 starts per rolling-origin fit for the ablation and 4 starts for the main recovery analyses.

Rescorla–Wagner (RW):

qₜ = qₜ₋₁ + α × (yₜ - qₜ₋₁), q₀ = 0.5

pₜ = qₜ₋₁ (clipped to [1e-9, 1-1e-9])

generation uses a fixed representative α = 0.30; the fitted α is free and is estimated by a coarse grid (49 points) plus L-BFGS-B refinement.

Fixed-gain restricted adaptive filter (λ identically 1): this competitor is the proposed four-parameter filter with the learning gain fixed at one (observation-precision increment π = π^ + 1.0 × s′(v)², i.e. λ is identically 1 for the learner), while the initial mean μ, initial σ and precision-decay ω are free. The original fitting box was μ in [-8, 8], log(ω) in [-8, 4], log(σ) in [-6, 6] (decision-variable order μ, log ω, log σ); a common-bounds sensitivity fits these shared parameters over μ in [-6, 6], log(ω) in [-4, 1], log(σ) in [-3, 2.5], matching the proposed model. The original routine returned fitted parameters with σ and ω swapped (a return-order bug); we corrected this, with details and a minimal numerical test in Supplementary Text S7. This is a restricted adaptive filter, not a standard Kalman filter: for Bernoulli–logistic observations the Kalman gain is not constant and does not match the form above. The internal code key kalman_fixed is kept only for reproducibility.

Competitor parity. In each comparison, every dataset was generated once and scored by all four models. Under the corrected protocol, the fixed-gain and proposed models use identical optimization (L-BFGS-B; same starts per prefix, deterministic initial draws, tolerances, and likelihood—fixed-gain is the proposed recursion with λ=1); the RW model uses grid plus refinement. All models had the same access to the training prefix in rolling-origin evaluation, and none saw held-out outcomes during fitting. We report pre-correction and corrected protocols separately (Table S18, Text S7); the corrected fit changes only the return order and, for sensitivity, the bounds.

## S2 Text. Mechanism-level decomposition of the perturbations

The main text groups perturbation analyses under robustness. To avoid conflating three distinct questions, they are separated as follows:

1. Recovery under the correctly specified model — main text, Parameter recovery, Identifiability, and Supplementary Table S7: data are generated and fit with the same model; this addresses identifiability.
2. Robustness to misspecification—data generated by alternative processes (RW, fixed-gain restricted filter) but fit with the proposed model: covered by model-recovery and ablation analyses (Model recovery, Model specification, Tables S8–S14). Poor recovery in this setting does not imply intrinsic non-identifiability.
3. Robustness to missing/corrupted observations: measurement noise perturbs observed binary outcomes; dropped responses remove a fraction of observations. The estimator is applied to the observed sequence as-is, without being told a perturbation occurred. These are secondary checks and do not support the identifiability conclusions.

## S3 Text. Terminology

Confidence-precision state (σ). σ is an internal confidence/precision-like state defined by the update equations; it is neither the inverse variance of a sampled Gaussian prior nor an empirically validated measure of psychological confidence. Thus, a fitted σ becoming large indicates weak information in this parameter under the chosen update equations, not an abnormal precision mechanism.

Fixed-gain restricted adaptive filter (code key kalman_fixed). The competitor with learning gain fixed at λ = 1 is a restricted form of the proposed adaptive filter (observation-precision increment fixed at s′(v)²), not a standard Kalman filter, whose gain for Bernoulli–logistic observations is not constant. The code key kalman_fixed is kept only for reproducibility and is not used in the manuscript.

Tolerance-based recovery rate: the proportion of replications in which the point estimate falls within a prespecified tolerance of the true parameter — here |θ^ - θ*| ≤ 0.6 on the log scale (positive parameters) or log-odds scale (μ). 0.6 log units corresponds to a factor of about 1.8 in a positive parameter or an odds ratio of about 1.8 for μ; it is a generous band relative to the errors documented here, not a universal success criterion. Sensitivity at 0.3 and 1.0 is reported in Table S9 (with the grid λ rates 0.078/0.182/0.263 distinguished from σ/ω, which stay at most 0.002); the conclusion does not hinge on the single 0.6 value.

Recovery check (likelihood audit) is correctly specified. The Table S7 experiment simulates and fits data under the same implemented model to test whether the boundary solution arises from the model’s own dynamics. This avoids misspecification from fitting a different model, but it does not show the algorithm is a correct model of human social learning, identify the source of remaining estimation issues, or by itself prove the implementation is correct—hence the independent-likelihood cross-check. These tests target the likelihood layer (recursion, sequence likelihood, initial-state convention, and finite-difference gradients) and are distinct from the Supplementary Text S6 evaluation-protocol tests, which ensure no forecast block is fit on the outcomes it is scored on.

Interval coverage/width describe how often uncertainty intervals contain the true parameter and how wide they are. We do not report these because many fits hit parameter boundaries, causing bootstrap/profile intervals to degenerate. Instead, we document the profile geometry (Identifiability Fig 1, S1 Fig, Table S17). Per the agreed scope, calibrated coverage is not assessed. The tolerance-based recovery rate is a point-estimate diagnostic, not interval coverage; neither flat profiles nor boundary estimates imply coverage. Wilson score intervals for recovery proportions and paired-bootstrap intervals for mean log-loss differences (Tables S11–S14) quantify uncertainty in proportions and paired means, not parameter-recovery coverage.

## S4 Text. Analysis-to-output map

**S1 Table.** Analysis-to-output map: main-text results, generating scripts, and archived outputs.

| Main-text result / element | Generating script | Archived output |
|---|---|---|
| R100 grid and trial-length recovery; bias/MAE/RMSE/boundary (Sec 3.1, Table S5) | review3_supplement.py | review3_diagnostics.json |
| Multi-tolerance recovery (0.3/0.6/1.0) and ranking (Sec 3.1, Table S9) | review3_supplement5.py | review3_diagnostics5.json |
| Three-observation model-recovery pilot; confusion matrix (Sec 3.3, Table S11) | review3_supplement.py / review3_supplement5.py | review3_diagnostics.json / review3_diagnostics5.json |
| Block model recovery (pre-correction), independent-likelihood cross-check, and second-regime profile (Tables S7/S17, Table S11) | review3_supplement7.py | review3_diagnostics7.json |
| Forecast-index validation and no-look-ahead causality test (Text S6) | review3_supplement9.py | review3_diagnostics9.json |
| Fixed-gain fitting parity: return-order correction and common-bounds sensitivity, corrected block model recovery and calibrated null (Sec 3.3/4.4, Tables S14/S18, Texts S6/S7) | review3_supplement10.py | review3_diagnostics10.json |
| Likelihood audit; NLL true vs fitted (Sec 3.1, Table S7) | review3_supplement3.py | review3_diagnostics3.json |
| Profile and trade-off diagnostics (Sec 3.5, Figs S1–S2) | supplement_figs.py | review3_diagnostics.json |
| Recovery scatterplots (Figs S3–S4) | make_fig_s3_s4.py | review3_diagnostics4.json |
| Authoritative paired ablation (single source; Sec 2.1, Tables S8/S12, Figs S5–S6); scatter/paired-difference raw arrays | review3_supplement8.py (tables) / review3_supplement4.py / make_fig_s5_s6.py (figures) | review3_diagnostics8.json (tables; supersedes earlier rounded ablation values) / review3_diagnostics4.json (figures) |
| Out-of-sample paired log-loss and bootstrap CIs (Sec 3.6, Table S13) | review3_supplement6.py | review3_diagnostics6.json |

Each main-text numerical result is linked to the script that generates it and the archived output it uses; both are available in the public repository (https://github.com/joevazhong-arch/practical-identifiability-adaptive-learning-gain-binary-feedback) described in Reproducibility of the main text. The manuscript version (V1.0.1) is archived on Zenodo with a persistent DOI (https://doi.org/10.5281/zenodo.23531449).

Dependencies are pinned, and the authors verified the full pipeline in a fresh virtual environment: ‘python reproduce.py’ regenerates all ten output families and six figures, and ‘python verify_reproduction.py’ matches every value to the archive within 1e-6 (28,972 values; max abs diff 0), with figures reproduced byte-for-byte (identical SHA-256). The package is released as a versioned archive: https://github.com/joevazhong-arch/practical-identifiability-adaptive-learning-gain-binary-feedback, V1.0.1 archived on Zenodo (https://doi.org/10.5281/zenodo.23531449). An independent clean-environment run by a non-author also reproduced the key results and matched the archive within tolerance. Together, these checks confirm the package regenerates the reported results.

## S5 Text. Profile geometry across regimes (MC1 cross-check)

S1 Fig profiles were first computed for one high-gain case (θ*=(0,3,0.02,5), T=120). To address the concern that the flat geometry might be scenario-specific, we recomputed profiles for a second regime with different boundaries, θ=(−1,1.5,0.02,2), T=120 (Table S17). In this regime, the MLE again pushed σ to its upper bound and λ to its lower bound (NLL 81.156). Across feasible ranges, absolute ΔNLL was 11.86 for μ, 0.26 for σ, 0.27 for ω (<1 NLL unit), and 9.39 for λ. The λ profile was flat for log λ ≤ 0 (ΔNLL ≤ 0.08; λ ≤ 1) and increased only for log λ ≥ 2, despite the true λ=2. This distinguishes likelihood information (profile geometry) from optimization performance (three-optimizer and dispersed-start checks). Thus, the flat σ–ω geometry and low-λ flatness replicate in a second regime, but this is not a claim over the full 24-condition grid: λ steepness (9.4 here vs 104.06 at high gain) and the σ–ω trade-off shape are regime-dependent and not generalized across parameter vectors.

## S6 Text. Forecast-index validation, no-look-ahead test and calibrated model-selection null

Validation uses two separate layers. Table S7 likelihood-layer checks (probability recursion, sequence likelihood, initial-state convention, finite-difference gradients, and an independent likelihood implementation) verify the likelihood code; the evaluation-protocol checks ensure rolling-origin scoring never fits on data it also scores.

**S2 Table.** Evaluation-protocol windows: non-overlapping fit/forecast splits.

| **Origin (cut)** | **Fit prefix (1-based)** | **Forecast block (1-based)** | **Scored trials** |
|---|---|---|---|
| t=24 | y1–y24 | y25–y44 | 20 |
| t=44 | y1–y44 | y45–y64 | 20 |
| t=64 | y1–y64 | y65–y80 | 16 |
| Total | — | union of the three blocks | 56 |

Evaluation uses non-overlapping windows. Parameters are fixed from the prefix fit, and the latent state is updated online using only past outcomes; thus the one-step-ahead probability pₜ for scoring yₜ is computed before observing yₜ and contains no information from it. Automated checks in review3_supplement9.py verify that, for every origin, fit and score index sets are disjoint (True), each future trial is scored exactly once (True; 56 distinct scored outcomes per dataset), and the final trial y80 is scored (True). Trials scored as “future” at earlier origins can later appear in the prefix at later origins (standard rolling-origin reuse), but are never used to fit the same origin in which they are scored. This is rolling-origin sequential prediction with prefix-fixed parameters: later predictions within a block may use earlier outcomes from that block, but never the outcome being scored or any future outcome.

No-look-ahead causality test: flipping one scored observation yₜ (trial 31) keeps pₜ and all earlier predictions unchanged, and changes only pₜ₊₁. The adaptive filter, Rescorla– Wagner, and the fixed-gain restricted filter pass (causality pass = True). Fit/score calls for dataset 1 are saved in review3_diagnostics9.json as an executable protocol trace.

**S3 Table.** Calibrated model-selection null: observed vs expected accuracy.

| **True model (common)** | **Marginal winner count (freq)** | **Observed correct / n** | **Null expected correct** | **Permutation p** |
|---|---|---|---|---|
| Adaptive filter | 26 (0.17) | 9/50 | 8.67 | 0.527 |
| Rescorla–Wagner | 80 (0.53) | 45/50 | 26.67 | <.001 |
| Fixed-gain restricted | 44 (0.29) | 18/50 | 14.67 | 0.140 |
| Overall | — | 72/150 | null mean 0.333 (SD 0.037) | <.001 |

Calibrated model-selection null (corrected). Under the corrected common-bounds protocol, selection is asymmetric: adaptive filter 26 (0.17), Rescorla–Wagner 80 (0.53), fixed-gain filter 44 (0.29) (corrected wide: 28/92/30; pre-correction: 54/84/12). The calibrated permutation null keeps per-dataset scores and winner labels fixed and permutes only generator labels (50,000 permutations; seed 20241018), preserving marginal winner frequencies. The table reports, for common-bounds, observed row-wise correct counts vs per-row null expectations and permutation p-values; the overall test is the sole inferential result.

Under the calibrated null, common-bounds accuracy is 0.333 (SD 0.037; 95% reference interval 0.260–0.407). The observed 72/150 = 0.480 gives permutation p < 0.001; the corrected wide protocol gives 67/150 = 0.447, p = 0.001; pre-correction 58/150 = 0.387 gave p = 0.079. Thus the corrected overall diagonal contains generator information but is strongly asymmetric: above-chance accuracy comes from the RW row (per-row p < 0.001), while the adaptive (0.527 common; 0.301 wide) and fixed-gain rows (0.140 common; 0.255 wide) do not differ from null. The overall test is the only inferential result; per-row values are descriptive, uncorrected, and not used to claim discrimination. The pre-correction fixed-gain 0/50 was due to the return-order defect (Text S7); corrected 12/50 or 18/50 remains < 0.5 and non-significant. A non-significant nested-pair result does not imply equivalence (which would require a pre-specified margin); the failure to separate the nested pair is specific to this evaluation procedure, not a general inability to distinguish model classes. The three-observation pilot (Table S11) lacks a permutation null because only aggregate scores (not per-dataset forecasts) were archived, and is kept only as a low-information reference.

## S7 Text. Fixed-gain fitting parity: return-order correction and common-bounds sensitivity

We ran a fitting-parity audit to address the reviewer’s concern that the fixed-gain restricted filter and the proposed model were not fit under comparable conditions. We identified two distinct discrepancies and treat them separately.

1. Return-order defect. The fixed-gain routine optimized z = (μ, log ω, log σ) and correctly evaluated θ = (μ, σ, ω), but returned (μ, ω, σ), swapping σ and ω. Since forward/simulate unpack (μ, σ, ω), forecasts were evaluated with exchanged parameters. A minimal test (test_kalman_order.py, seed=7) yields NLL 50.4632; the defective path gives 1493.1953; restoring (μ, σ, ω) returns 50.4632. competitors.py was fixed to return (bestz[0], exp(bestz[2]), exp(bestz[1])) = (μ, σ, ω); the objective was already correct.
2. Fitting bounds. The fixed-gain model used μ∈[-8,8], log ω∈[-8,4], and log σ∈[-6,6] (order: μ , log ω , log σ ). The proposed model uses μ∈[-6,6], log σ∈[-3,2.5], log ω∈[-4,1], and log_lambda∈[-4,4]. For common-bounds sensitivity, shared fixed-gain parameters are fit over μ∈[-6,6], log ω∈[-4,1], and log σ∈[-3,2.5].

Comparable effort. Under the corrected protocols, both models use the same L-BFGS-B optimizer, n_starts=2 per prefix, deterministic starting draws (seed=0), convergence tolerances, and likelihood (forward_kalman uses the proposed forward recursion with λ=1), making likelihoods and convergence comparable; the only differences are return order and bounds. All fixed-gain comparisons in the main text and supplement use the corrected routine; pre-correction figures appear only as a clearly labelled historical control.

Separated effects and conclusion. Correcting the return order moves the overall accuracy 58→67 (and changes 38 winners); moving to common bounds moves it 67→72 (a further 26 winners). The corrected overall accuracy is modestly above chance (0.333), with a significant calibrated permutation test (p = 0.0011 wide; p < 0.001 common), but this is driven by the RW row (0.88–0.90); the adaptive and fixed-gain rows (0.18–0.22 and 0.24–0.36) remain non-significant and mutually confused. The main claim—that the learning gain λ cannot be reliably distinguished from its fixed-gain restriction—therefore survives and is reported as a failure to discriminate the nested pair under the tested evaluation procedure, not as a general inability to distinguish model classes; it does not prove the restricted model false or the full adaptive mechanism correct, and it is robust to the fitting bounds.

## S8 Text. Recovery-practice audit of representative BPD computational-psychiatry studies

**S4 Table.** Recovery-practice audit of six representative BPD computational-psychiatry studies.

| Study | Recovery simulation? | Reported recovery | Primary claim level |
|---|---|---|---|
| **Siegel 2020 [9]** | No | n/a | Single-parameter group difference (belief certainty toward bad agents) |
| **Henco 2020 [10]** | Yes (estimate→estimate) | Weak; ω<sub>3</sub>,cue recovery flagged as poor | Single-parameter group difference |
| **Fineberg 2018 [12]** | No | n/a | Single-parameter group difference (reversal learning) |
| **Csukly 2023 [13]** | No (only MCMC diagnostics) | n/a | Single-parameter group difference (well-being weights) |
| **Story 2024 [14]** | Yes | ψ_Bad r=0.64; ψ_Split r=0.74 | Single-parameter group difference (splitting) |
| **Barnby 2025 [11]** | Yes (full) | r=0.70–0.94 | Model-structural difference (self–other separation) |

*Note: “Yes” means the authors ran a simulation-based parameter-recovery check on their task; “Weak” means at least one clinically central parameter had a recovery coefficient below r=0.7.*

We reviewed the Methods of six representative BPD computational-psychiatry studies using Bayesian or reinforcement-learning parameter estimation in social feedback tasks. For each study, we noted whether authors performed simulation-based parameter recovery for their task, reported any recovery coefficient, and the scope of the primary claim (single-parameter group difference, model-structural difference, or predictive comparison). This descriptive audit is not a quality judgment; it contextualizes the recovery standard proposed in the Implications for empirical BPD computational psychiatry section.

**S5 Table.** Recovery metrics for Experiment 1 (parameter grid, fixed T=80)

| **Parameter** | **Signed bias** | **MAE** | **RMSE** | **Tol. recovery (≤0.6)** | **Boundary hits / fit (of 4)** |
|---|---|---|---|---|---|
| μ | -0.09 | 2.49 | 2.78 | 0.06 | 0.41 |
| σ | +2.57 | 3.25 | 3.30 | 0.001 | 3.71 |
| ω | +0.18 | 2.00 | 2.20 | 0.002 | 3.53 |
| λ | -1.71 | 2.49 | 3.22 | 0.18 | 1.53 |

Experiment 1: 24-condition parameter grid (vary generating parameters; fixed T=80; 100 replications per condition; 4 deterministic starts per fit). Aggregated across conditions; μ on the log-odds scale; σ/ω/λ on the log scale.

**S6 Table.** Recovery metrics for Experiment 2 (trial-length analysis)

| Parameter | Signed bias (range) | MAE (range) | RMSE (range) | Tol. recovery (range) | Boundary hits / fit (range) |
|---|---|---|---|---|---|
| μ | -0.14 … +0.07 | 0.67–0.92 | 0.82–1.14 | 0.38–0.55 | 0 |
| σ | +1.807 (constant) | 1.807 | 1.807 | 0 | 3.88–3.95 |
| ω | +0.49 … +3.96 | 1.89–3.96 | 2.33–3.96 | 0 | 2.80–3.80 |
| λ | -4.46 … -4.19 | 4.19–4.46 | 4.22–4.59 | 0 (0.01 at T=80) | 0.25–2.79 |

Experiment 2: Trial-length analysis using representative vector *θ* \*=(0.5,2.0,0.05,3.0). T ∈ {40, 80, 120, 240, 480}, with 100 replications per T. Ranges are reported across the five trial counts.
Interpretation. The designs differ: Experiment 1 varies generating parameters at a fixed trial count, while Experiment 2 varies trial count for one representative parameter vector. μ performs better in Experiment 2 (MAE 0.67–0.92) but not across Experiment 1’s full grid (MAE 1.9–3.2), so μ is not reliably recovered.

**S7 Table.** Correctly specified recovery check (likelihood audit)

| **Quantity** | **Mean** | **SD** | **95% CI** |
|---|---|---|---|
| NLL at true vector | 45.23 | 10.49 | 42.33–48.14 |
| NLL at fitted optimum | 43.65 | 10.57 | 40.72–46.58 |
| Difference (fitted minus true) | -1.59 | 0.86 | -1.82 … -1.35 |

*θ* \*=(0.5,2.0,0.05,3.0), T=80, R=50 independent replications, 4 deterministic starts; the true vector is evaluated using the same forward recursion, initial-state convention, and likelihood indexing as the fitted objective (review3\_diagnostics3.json). This correctly specified recovery check tests whether the boundary solution can arise under the data-generating dynamics, not whether the implementation is formally proven correct.

The fitted optimum outperformed the true vector in 50/50 replications. Boundary-hit rates: σ 1.00, ω 0.96, λ 0.48, μ 0. Because data were generated and fit with the same implemented model, this shows boundary solutions can occur even when the fitted model matches the generator, easing concern about some implementation errors. However, it does not show the algorithm is a correct model of human social learning or explain the boundary solution. Nor does it prove the likelihood code is correct: shared simulator/evaluator bugs could still yield agreement.

Independent likelihood cross-check. To avoid shared implementation errors, we re-implemented the forward recursion as a plain scalar loop in pure Python (math.exp only; no numpy; distinct variable names; no imports from adaptive_learning_gain.py). Across 150 random sequences spanning model-recovery regimes, both versions matched with max absolute one-step probability difference 0.000e+00 and max absolute NLL difference 8.527e-14 (tolerances 1e-9 and 1e-6; pass=true). The independent recursion also reproduces the worked-example first three probabilities (0.622459, 0.656197, 0.684670) and yields a finite NLL for an extreme 50-ones/50-zeros sequence (201.8096), matching the main implementation. Along with same-codebase unit tests (manual five-trial NLL and finite-difference gradient check), we find no issues in recursion, indexing, or initial-state conventions; however, this is not a formal proof, since any shared error would evade both checks.

**S8 Table.** Paired ablation comparison on identical shared datasets.

| Experiment / setting / trial length | Comparison | Mean diff ± SD | Normal 95% CI for mean | Full better (datasets) |
|---|---|---|---|---|
| M1 (T=80) moderate (0.5,2,.05,3) | full vs no-s' | -0.00442 ± 0.05626 | -0.02456 to +0.01571 | 17/30 |
| M1 (T=80) moderate (0.5,2,.05,3) | full vs simpler-precision | +0.02010 ± 0.13177 | -0.02706 to +0.06725 | 13/30 |
| M1 (T=80) moderate (0.5,2,.05,3) | full vs RW | -0.02296 ± 0.13136 | -0.06997 to +0.02404 | 19/30 |
| M1 (T=80) high-gain (0,3,.1,5) | full vs no-s' | -0.01312 ± 0.07895 | -0.04138 to +0.01513 | 11/30 |
| M1 (T=80) high-gain (0,3,.1,5) | full vs simpler-precision | +0.01239 ± 0.06305 | -0.01017 to +0.03495 | 20/30 |
| M1 (T=80) high-gain (0,3,.1,5) | full vs RW | -0.02471 ± 0.17585 | -0.08764 to +0.03821 | 14/30 |
| M1 (T=80) low-gain (-1,1.5,.02,2) | full vs no-s' | -0.02875 ± 0.07152 | -0.05435 to -0.00316 | 17/30 |
| M1 (T=80) low-gain (-1,1.5,.02,2) | full vs simpler-precision | +0.01430 ± 0.05192 | -0.00428 to +0.03288 | 16/30 |
| M1 (T=80) low-gain (-1,1.5,.02,2) | full vs RW | -0.07494 ± 0.18738 | -0.14199 to -0.00788 | 21/30 |
| M2 (T=40) moderate | full vs no-s' | -0.01644 ± 0.09305 | -0.06354 to +0.03065 | 6/15 |
| M2 (T=40) moderate | full vs simpler-precision | -0.00471 ± 0.01296 | -0.01127 to +0.00184 | 10/15 |
| M2 (T=40) moderate | full vs RW | +0.01372 ± 0.12241 | -0.04823 to +0.07567 | 7/15 |
| M2 (T=80) moderate | full vs no-s' | -0.00993 ± 0.04639 | -0.03341 to +0.01354 | 6/15 |
| M2 (T=80) moderate | full vs simpler-precision | -0.03047 ± 0.07282 | -0.06732 to +0.00639 | 11/15 |
| M2 (T=80) moderate | full vs RW | +0.01904 ± 0.06381 | -0.01325 to +0.05134 | 5/15 |
| M2 (T=120) moderate | full vs no-s' | +0.00130 ± 0.00392 | -0.00068 to +0.00329 | 5/15 |
| M2 (T=120) moderate | full vs simpler-precision | -0.00460 ± 0.01053 | -0.00993 to +0.00074 | 11/15 |
| M2 (T=120) moderate | full vs RW | +0.01402 ± 0.05377 | -0.01319 to +0.04123 | 5/15 |

Two experiments are reported separately to prevent confusion from shared T=80 entries. M1 uses three generating settings (moderate, high-gain, low-gain) at fixed T=80. M2 uses one moderate setting (0.5,2,0.05,3) at T=40, 80, and 120. Each dataset was generated once with the full model, then scored by all four models using the same rolling-origin cuts (0.25/0.40/0.60/0.80). We report, per dataset, the full-minus-variant difference in mean held-out log-loss; negative values favor the full model. All results are recomputed from the authoritative full-precision per-dataset array (review3_diagnostics8.json; five decimals), match Table S12, and replace earlier rounded values, removing the previous one-count discrepancy from win tallies based on rounded differences.

**S9 Table.** Tolerance-based recovery rates (Experiment 1 grid, T=80)

| **Parameter** | **Tol ≤ 0.3** | **Tol ≤ 0.6** | **Tol ≤ 1.0** |
|---|---|---|---|
| μ | 0.032 | 0.062 | 0.107 |
| σ | 0.001 | 0.001 | 0.002 |
| ω | 0.000 | 0.002 | 0.002 |
| λ | 0.078 | 0.182 | 0.263 |

Part A. Tolerance-based recovery rates for Experiment 1 (T=80), pooled across 24 conditions and 100 replications (n=2400), at three transformed-scale tolerances (μ: log-odds; σ/ω/λ: log). Results show the qualitative conclusion is not driven by the 0.6 threshold (review3_diagnostics5.json).

Experiment 1 shows parameter-specific recovery: σ and ω are essentially unrecoverable (≤0.002 at all tolerances), while λ has low but non-negligible recovery that increases with tolerance (0.078/0.182/0.263 at 0.3/0.6/1.0). The experiments should not be merged: in Experiment 2 (single parameter vector), λ shows almost no recovery (0–0.01), which does not generalise to the full 24-condition grid. In Experiment 2, μ recovery at tolerances 0.3/0.6/1.0 is 0.29/0.51/0.66 at T=80, rising to ∼0.74 by T=120 and longer trials; σ and ω remain ∼0 across tolerances and trial lengths. Even at tolerance 1.0 (≈2.7× in original units), σ and ω stay ≤0.002 across the grid, whereas λ is not near-zero.

**S10 Table.** Rank-based recovery (Spearman ρ between true and estimated parameters)

| **Parameter** | **Pooled Spearman ρ (n=2400)** | **Condition-mean Spearman ρ (n=24)** |
|---|---|---|
| μ | 0.306 (p<.001) | 0.867 (p<.001) |
| σ | 0.063 (p=0.002) | 0.199 (p=0.352) |
| ω | 0.004 (p=0.850) | 0.096 (p=0.654) |
| λ | 0.242 (p<.001) | 0.907 (p<.001) |

Part B. Rank-based recovery: Spearman rank correlation between true and estimated transformed parameters at two aggregation levels. Pooled uses all n=2400 individual estimates; condition-mean uses n=24 condition means (one per generating vector). p-values are asymptotic two-sided. Pooled reflects individual ranking; condition-mean reflects easier across-regime (ecological) ordering. The trial-length experiment uses one generating vector and cannot assess ranking.

Ranking of individual sequences is weak even for μ (pooled ρ 0.31) and absent for σ and ω (p non-significant); the high condition-mean ρ for μ and λ reflects only ordering of the generating regimes, not ordering of individuals within a regime, which is the level at which phenotyping would operate.

**S11 Table.** Model-recovery pilot: three-observation rolling-origin scoring.

| **Generating model** | **Correct / n** | **Accuracy** | **Wilson 95% CI** | **Reference** |
|---|---|---|---|---|
| Overall | 57/150 | 0.38 | 0.306–0.460 | chance 0.333 (inside) |
| Adaptive filter | 13/50 | 0.26 | 0.159–0.396 | — |
| Rescorla–Wagner | 31/50 | 0.62 | 0.481–0.741 | — |
| Fixed-gain restricted filter | 13/50 | 0.26 | 0.159–0.396 | — |

Low-information pilot: each candidate is fit to three prefix cuts (t = 24, 44, 64) and scored only on the next trial yₜ₊₁, yielding just three binary forecast observations per dataset. We retain this pilot to assess sensitivity to the number of scored future observations; the primary block protocol scores 56 future trials per dataset (Table S14). This pilot is only a reference analysis, and we do not compute a calibrated permutation null because the archived output contains only aggregate scores, not per-dataset point forecasts; the calibrated permutation null is reported only for the primary block protocol (Text S6). We report Wilson 95% CIs for overall accuracy and row-wise recovery rates, and show the three-model chance level (0.333) for reference (review3_diagnostics5.json; review3_diagnostics7.json). The pilot overall CI (0.306–0.460) includes 0.333. The corrected block protocol, with ∼19× more forecast observations per dataset, increases overall accuracy only to 0.45–0.48 (Table S14), driven by the RW row; results are finite-sample (n=50 per row) and apply only to these three implementations and simulation settings.

**S12 Table.** Paired ablation intervals (normal and bootstrap)

| Parameter | Tol ≤ 0.3 |  | Tol ≤ 0.6 | Tol ≤ 1.0 |  |
|---|---|---|---|---|---|
| M1 moderate (T=80) | full vs no-s' | -0.00442 | -0.02456 to +0.01571 | -0.02427 to +0.01476 | 17/30 |
| M1 moderate (T=80) | full vs simpler-precision | +0.02010 | -0.02706 to +0.06725 | -0.02036 to +0.07073 | 13/30 |
| M1 moderate (T=80) | full vs RW | -0.02296 | -0.06997 to +0.02404 | -0.06685 to +0.02420 | 19/30 |
| M1 high-gain (T=80) | full vs no-s' | -0.01312 | -0.04138 to +0.01513 | -0.04339 to +0.01273 | 11/30 |
| M1 high-gain (T=80) | full vs simpler-precision | +0.01239 | -0.01017 to +0.03495 | -0.00599 to +0.03814 | 20/30 |
| M1 high-gain (T=80) | full vs RW | -0.02471 | -0.08764 to +0.03821 | -0.09273 to +0.02817 | 14/30 |
| M1 low-gain (T=80) | full vs no-s' | -0.02875 | -0.05435 to -0.00316 | -0.05649 to -0.00622 | 17/30 |
| M1 low-gain (T=80) | full vs simpler-precision | +0.01430 | -0.00428 to +0.03288 | -0.00122 to +0.03496 | 16/30 |
| M1 low-gain (T=80) | full vs RW | -0.07494 | -0.14199 to -0.00788 | -0.14643 to -0.01607 | 21/30 |
| M2 moderate (T=40) | full vs no-s' | -0.01644 | -0.06354 to +0.03065 | -0.06467 to +0.02366 | 6/15 |
| M2 moderate (T=40) | full vs simpler-precision | -0.00471 | -0.01127 to +0.00184 | -0.01188 to -0.00002 | 10/15 |
| M2 moderate (T=40) | full vs RW | +0.01372 | -0.04823 to +0.07567 | -0.04973 to +0.07037 | 7/15 |
| M2 moderate (T=80) | full vs no-s' | -0.00993 | -0.03341 to +0.01354 | -0.03335 to +0.01132 | 6/15 |
| M2 moderate (T=80) | full vs simpler-precision | -0.03047 | -0.06732 to +0.00639 | -0.07061 to +0.00032 | 11/15 |
| M2 moderate (T=80) | full vs RW | +0.01904 | -0.01325 to +0.05134 | -0.01322 to +0.04889 | 5/15 |
| M2 moderate (T=120) | full vs no-s' | +0.00130 | -0.00068 to +0.00329 | -0.00054 to +0.00327 | 5/15 |
| M2 moderate (T=120) | full vs simpler-precision | -0.00460 | -0.00993 to +0.00074 | -0.01036 to -0.00032 | 11/15 |
| M2 moderate (T=120) | full vs RW | +0.01402 | -0.01319 to +0.04123 | -0.01302 to +0.04004 | 5/15 |

For each experiment/setting/comparison, the table reports the mean paired full–variant difference, 95% CIs from a normal approximation and paired bootstrap (10,000 resamples of dataset-level differences), and the number of datasets where the full model had higher log-loss. Values are recomputed from the same full-precision per-dataset array as Table S8 (review3_diagnostics8.json, five decimals). A CI that excludes zero favours the full model only if both the mean and interval are negative.

Recomputed from the single authoritative array, all nine M2 normal-approximation intervals include zero. In M1, two of nine comparisons exclude zero under both normal and bootstrap intervals: low-gain full-vs-no-s ′ and full-vs-RW. In M2, under the bootstrap only the T=40 and T=120 full-vs-simpler-precision upper bounds are slightly negative (−0.00002, -0.00032); T=80 is +0.00032 and flips sign across resampling seeds (negative in 2/10). Because these effects are <0.005 log-loss per trial, disappear at normal precision, and are seed-sensitive, they are not treated as material or consistent. Earlier three-decimal rounding printed endpoints as -0.000/0.000 and produced one win-count mismatch with Table S8; all entries here use a single full-precision source, so Tables S8 and S12 now match exactly. Overall, the ablation suggests the added update components do not provide a consistent, material predictive benefit under these settings, though this does not prove they are never useful.

**S13 Table.** Out-of-sample paired comparison (50 datasets, fixed split)

| Paired comparison (model minus baseline) | Mean | SD | Normal 95% CI | Bootstrap 95% CI | Model better/worse |
|---|---|---|---|---|---|
| Adaptive model minus empirical-frequency | -0.024 | 0.111 | -0.055 to +0.007 | -0.058 to +0.003 | 60%/40% |
| Adaptive model minus constant s(μ̂) | -0.142 | 0.259 | -0.213 to -0.070 | -0.221 to -0.078 | 78%/22% |

R=50 independent datasets were generated from (0.5, 2.0, 0.05, 3.0) at T=120 using a fixed 72/48 train–test split; all three predictors were re-estimated for each training prefix. The empirical-frequency predictor used the raw training mean (MLE) and was held fixed on the test trials; frequencies were clipped to [1e-6, 1-1e-6] only for numerical stability (no Laplace/Bayesian shrinkage). Training frequencies ranged from 0.167 to 0.917 with no clipping.

Mean held-out log-loss: adaptive 0.520 (SD 0.202), empirical-frequency 0.544 (SD 0.208), constant s(μ^) 0.661 (SD 0.286). Data were generated by the proposed model, giving it a specification advantage, but finite-sample estimation error, flexibility, and the forecast setup can still prevent better held-out performance. No clear held-out gain over the empirical-frequency baseline was shown; this result depends on the tested generator, sample size, fitting procedure, and baseline and does not imply equivalence or general inferiority. The empirical-frequency predictor is only one constant-frequency baseline, not the full class of simpler sequential models. With intervals spanning zero and no pre-specified equivalence margin, this is a failure to show an advantage, not evidence of equivalence.

**S14 Table.** Corrected block model recovery (primary): 56 future trials per dataset.

| Protocol / generating model | Correct / n | Accuracy | Wilson 95% CI | Reference |
|---|---|---|---|---|
| Overall — common bounds (primary) | 72/150 | 0.480 | 0.402–0.559 | chance 0.333 (below) |
| Overall — corrected wide bounds | 67/150 | 0.447 | 0.369–0.527 | chance 0.333 (below) |
| Overall — pre-correction (control) | 58/150 | 0.387 | 0.312–0.467 | chance 0.333 (inside) |
| Adaptive filter (common) | 9/50 | 0.18 | 0.098–0.308 | — |
| Rescorla–Wagner (common) | 45/50 | 0.90 | 0.786–0.957 | — |
| Fixed-gain restricted (common) | 18/50 | 0.36 | 0.241–0.499 | — |

Primary rolling-origin model recovery (corrected). A fitting-parity audit (Supplementary Text S7) identified and fixed a return-order bug in the fixed-gain candidate, then re-fit shared parameters under bounds matching the proposed model. Each candidate is fit at three origins and scored (with parameters fixed) by one-step-ahead forecasts on the next non-overlapping block: fit y1–y24 → forecast y25–y44; fit y1–y44 → forecast y45–y64; fit y1–y64 → forecast y65–y80 (56 future trials total; each scored once, including y80).

R=50 datasets per generating model; all candidates use the same blocks. With corrected common bounds, overall accuracy is 72/150=0.480 (Wilson 0.401–0.559); with corrected but original wider bounds, 67/150=0.447 (0.369–0.527). The pre-correction protocol reproduces 58/150=0.387 and is kept as a historical control. Row-wise recovery is reported with Wilson 95% intervals (chance 0.333); indexing and no-look-ahead are enforced by automated assertions, and the calibrated permutation null is in Supplementary Text S6 (review3_supplement10.py, review3_diagnostics10.json).

**S15 Table.** Common-bounds block confusion matrix.

| **True \ Predicted (common)** | **Adaptive filter** | **Rescorla–Wagner** | **Fixed-gain restricted** |
|---|---|---|---|
| Adaptive filter | 9 | 18 | 23 |
| Rescorla–Wagner | 2 | 45 | 3 |
| Fixed-gain restricted | 15 | 17 | 18 |

Common-bounds block confusion matrix (rows = true generating model, columns = predicted winner; counts of 50 datasets).

**S16 Table.**
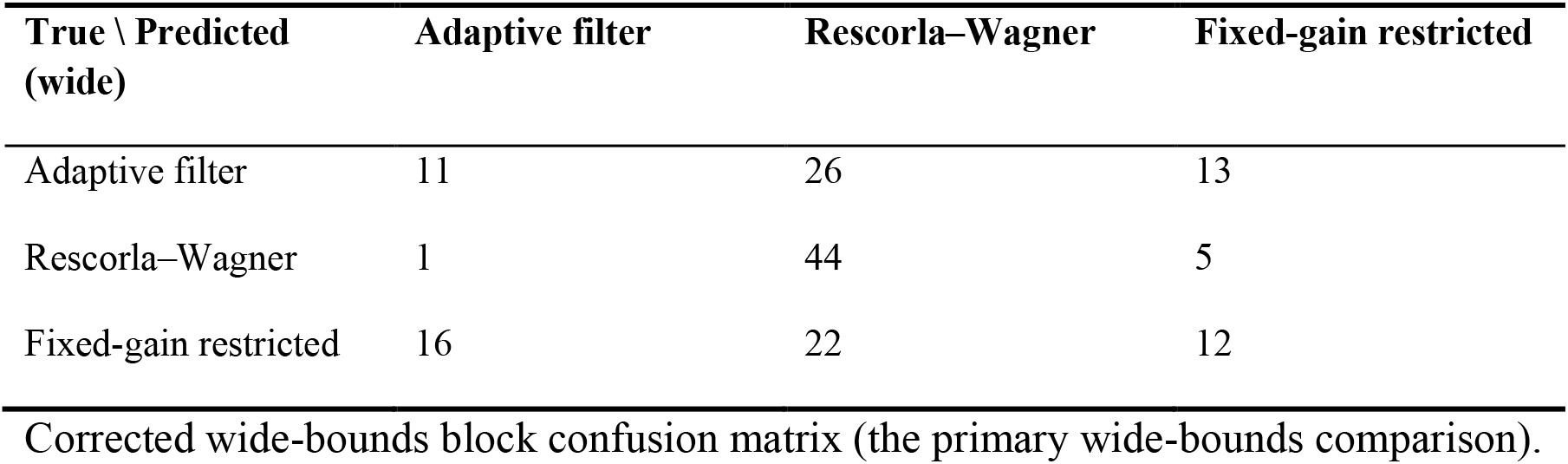
Corrected wide-bounds block confusion matrix.

| **True \ Predicted (wide)** | **Adaptive filter** | **Rescorla–Wagner** | **Fixed-gain restricted** |
|---|---|---|---|
| Adaptive filter | 11 | 26 | 13 |
| Rescorla–Wagner | 1 | 44 | 5 |
| Fixed-gain restricted | 16 | 22 | 12 |

Corrected wide-bounds block confusion matrix (the primary wide-bounds comparison).

After correction, the RW row is reliably identified (0.90 with common bounds; 0.88 with wide; per-row permutation p < 0.001). Adaptive and fixed-gain remain non-significant (0.18–0.22; 0.24–0.36) and are often confused: under common bounds, 23/50 adaptive datasets are labeled fixed-gain and 15/50 fixed-gain datasets are labeled adaptive. Fixed-gain (adaptive with λ = 1) is nested in the four-parameter model; its pre-correction 0/50 was due to the return-order defect (Text S7), and post-correction performance (12/50 wide; 18/50 common) is still < 0.5 and non-significant. Pre-correction winner–runner-up mean log-loss margins (SD) are small for all true models: adaptive -0.0401 (0.0469), RW -0.0244 (0.0483), fixed-gain -0.0221 (0.0205). Results are robust to bounds (wide→ common: 0.447→0.480; 26 winners changed); many wide-bound prefix estimates fall outside the common box ( σ : 435/450; ω : 328/450) without changing direction. These conclusions apply only to the tested implementations and simulation settings and do not establish equivalence (no pre-specified margin) or a general inability to distinguish model classes.

**S17 Table.**
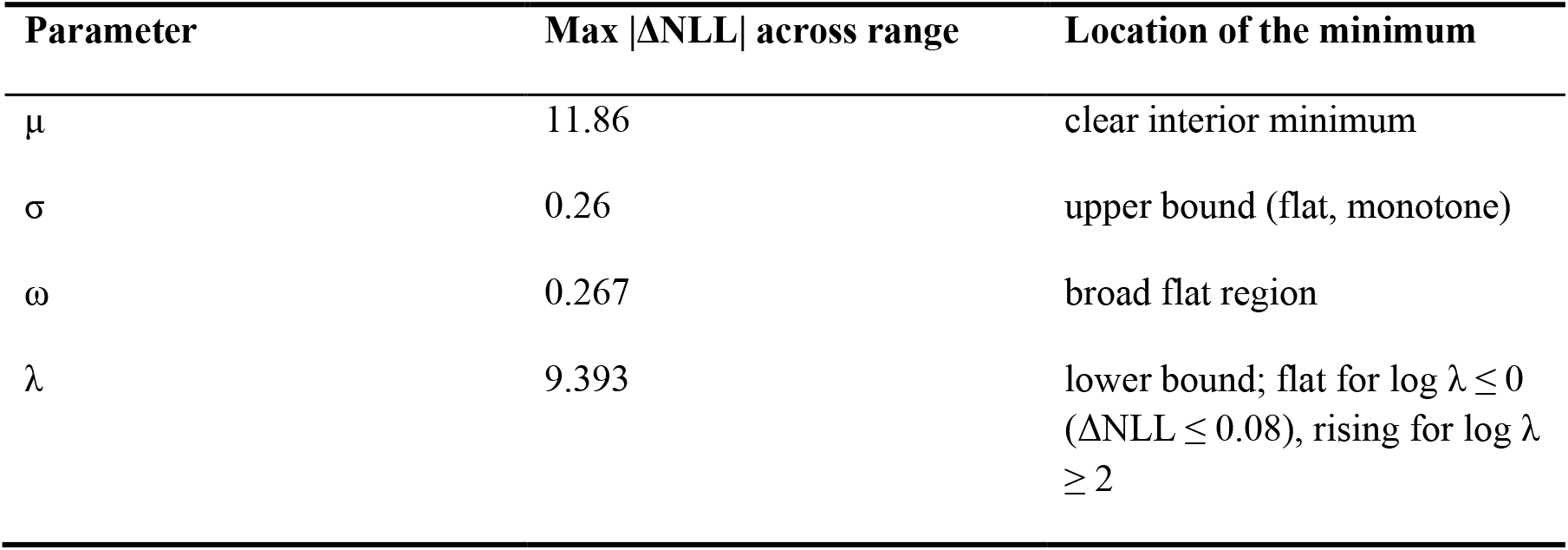
Profile-likelihood geometry in a second (low-gain) regime.

To check whether the flat profile geometry in S1 Fig is a high-gain artefact, we recomputed all four profiles under a low-gain regime with a different parameter vector and boundary pattern, θ*=(−1.0,1.5,0.02,2.0), T=120 (MLE NLL 81.156; the MLE again pushed σ to its upper bound and λ to its lower bound). The table reports the maximum absolute change in profile NLL over each parameter’s feasible range after re-optimising the others (review3_diagnostics7.json).

The flat σ–ω geometry (≤1 NLL change) and low-λ flatness also appear in this second regime, so they are not unique to high gain; however, this is shown only for two regimes, not the full 24-condition grid. The λ profile steepness is regime-dependent (span 9.39 here vs 104.06 at high gain), and μ is the only parameter with a clear minimum in both regimes. Thus, neither the profile shapes nor the σ–ω trade-off generalise across parameter vectors. See Supplementary Text S5.

**S18 Table.** Fixed-gain fitting parity: three-protocol comparison.

| **Protocol** | **Overall k/n** | **Accuracy** | **Wilson 95%** | **Permutation p (overall)** | **Fixed-gain row correct** | **Fixed-gain winners (of 150)** |
|---|---|---|---|---|---|---|
| Pre-correction (V15) | 58/150 | 0.387 | 0.312–0.467 | 0.079 | 0/50 | 12 |
| Corrected wide bounds | 67/150 | 0.447 | 0.369–0.527 | 0.001 | 12/50 | 30 |
| Corrected common bounds | 72/150 | 0.480 | 0.402–0.559 | <.001 | 18/50 | 44 |

Fitting-parity comparison of three protocols. The fixed-gain restricted filter is equation-nested (λ=1), but the original code differed from the proposed model in return order and fitting bounds. review3_supplement10.py evaluates three protocols on the same 150 datasets (same origins; 56 forecast trials; L-BFGS-B; n_starts=2 per prefix; identical deterministic starts, tolerances, and likelihood): (1) pre-correction (reproduces V15), (2) corrected fit with the original wider bounds, and (3) corrected fit with bounds matching the proposed model.

Boundary sensitivity with corrected wide bounds. In the 450 fixed-gain prefix fits (50 datasets × 3 origins), estimates often fall outside the common box: σ in 435, ω in 328, μ in 30, and at least one parameter in all 450. Switching to common bounds changes only magnitude, not direction, so the model-selection conclusion is robust to the bounds.

Separate effects on the winner: the return-order correction changes the winner on 38 datasets (58→67), and the bound change alters 26 more (67→72). Thus, return-order has the bigger effect, with bound change secondary; the three-protocol chain is 58→67→72.

## Supporting figures

**S1 Fig.**
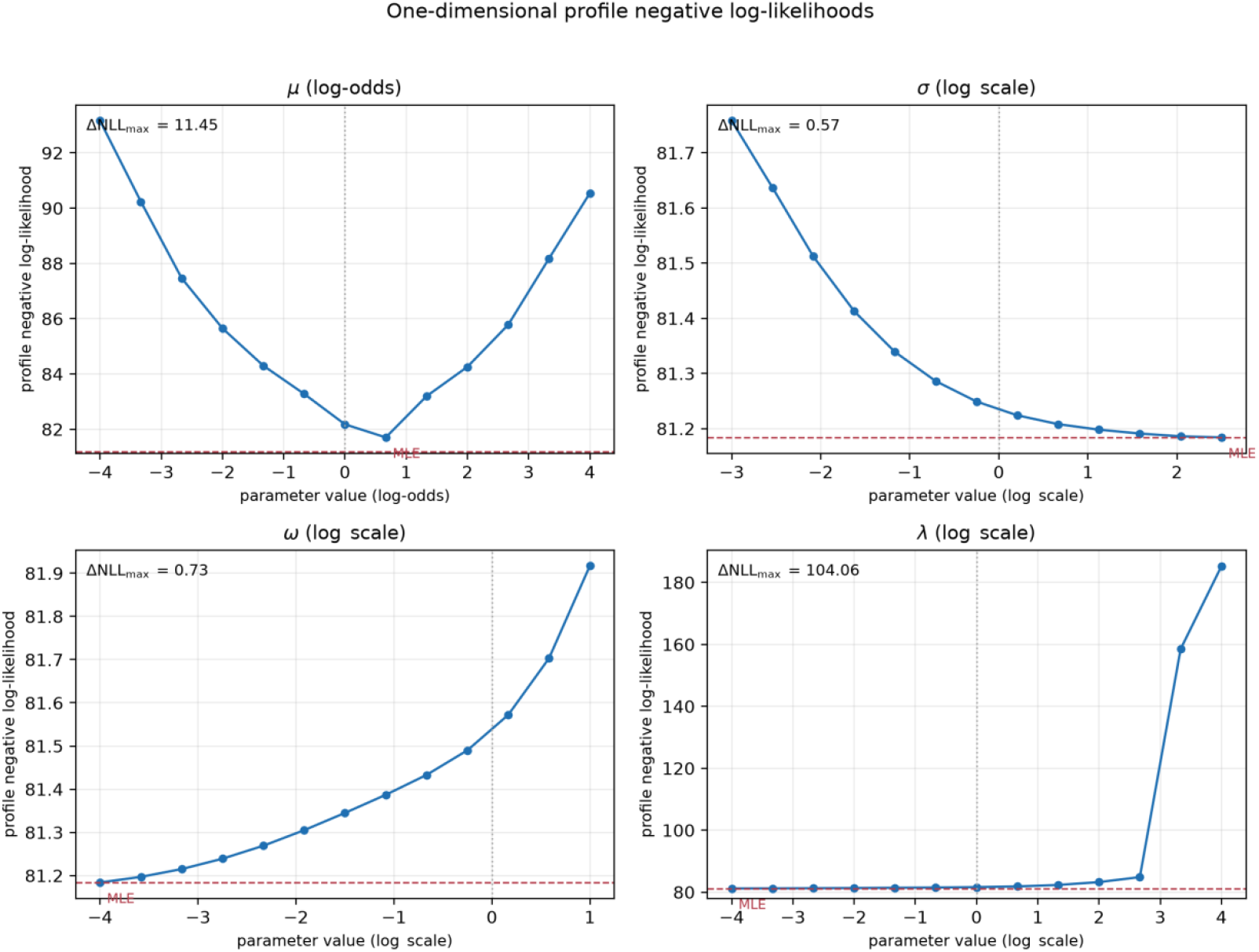
One-dimensional profile negative log-likelihoods for a representative high-gain scenario (θ*=(0,3,0.1,5), T=120; MLE NLL=81.184). The x-axis shows log-odds for μ and log values for σ, ω, and λ (bounded ranges); the y-axis shows profile NLL after re-optimising the other three parameters. The MLE line and the maximum NLL change for each profile are indicated. σ and ω are nearly flat (<1 NLL unit change); λ is flat below ∼2 but rises steeply at higher values (asymmetric); μ has a clear single minimum. The flat σ–ω geometry also appears in a second low-gain regime (Table S9) and only in 2 of the tested regimes; λ steepness varies by regime.

**S2 Fig.**
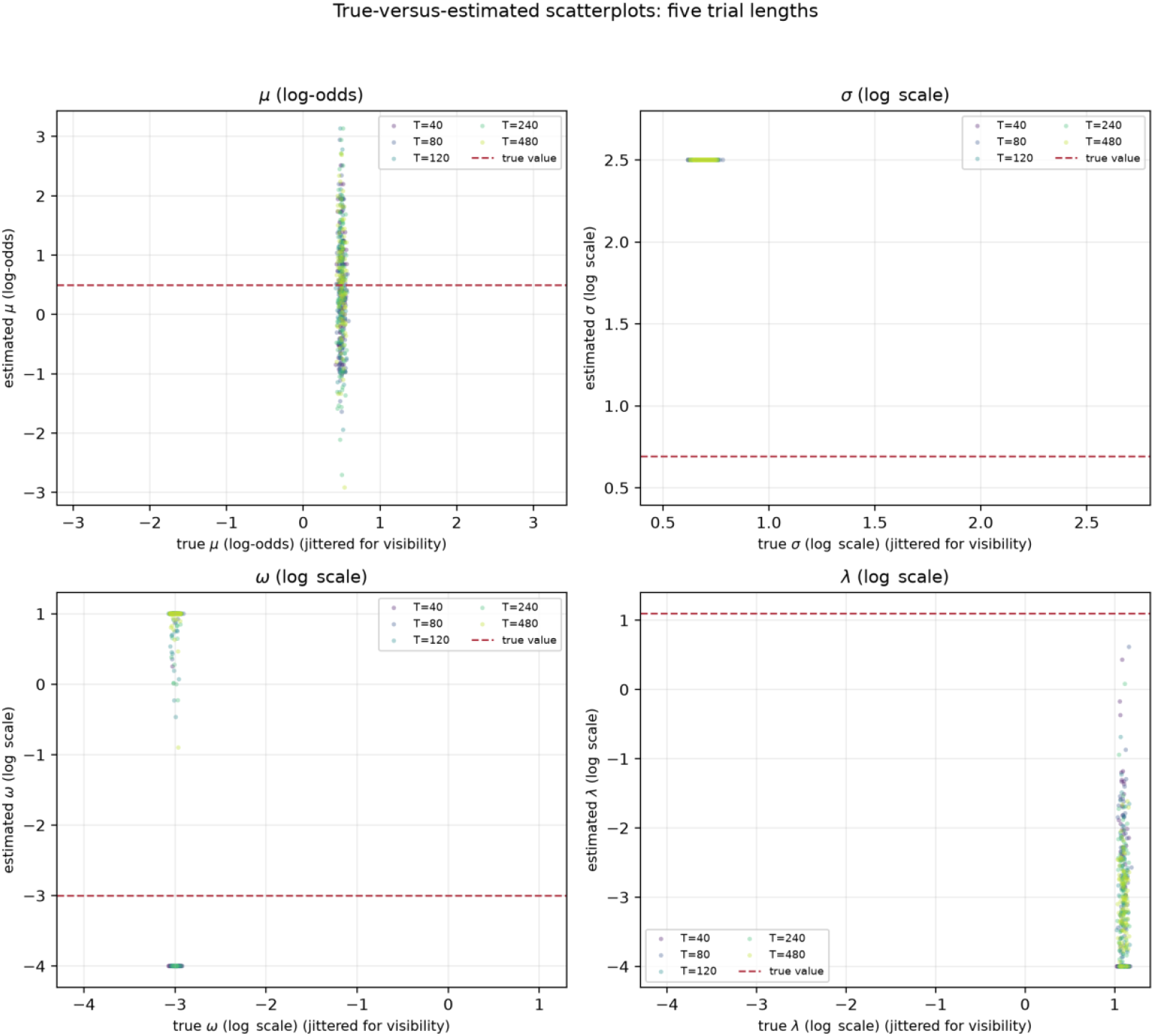
True vs. estimated scatterplots for Experiment 2 at θ *=(0.5,2.0,0.05,3.0) across trial lengths T∈ {40,80,120,240,480} (100 replications each). Axes use the same transformed scales as Fig 1A (log-odds for μ ; log for positive parameters); dashed red line shows the true value. Increasing T does not shift the positive parameters off their bounds.

**S3 Fig.**
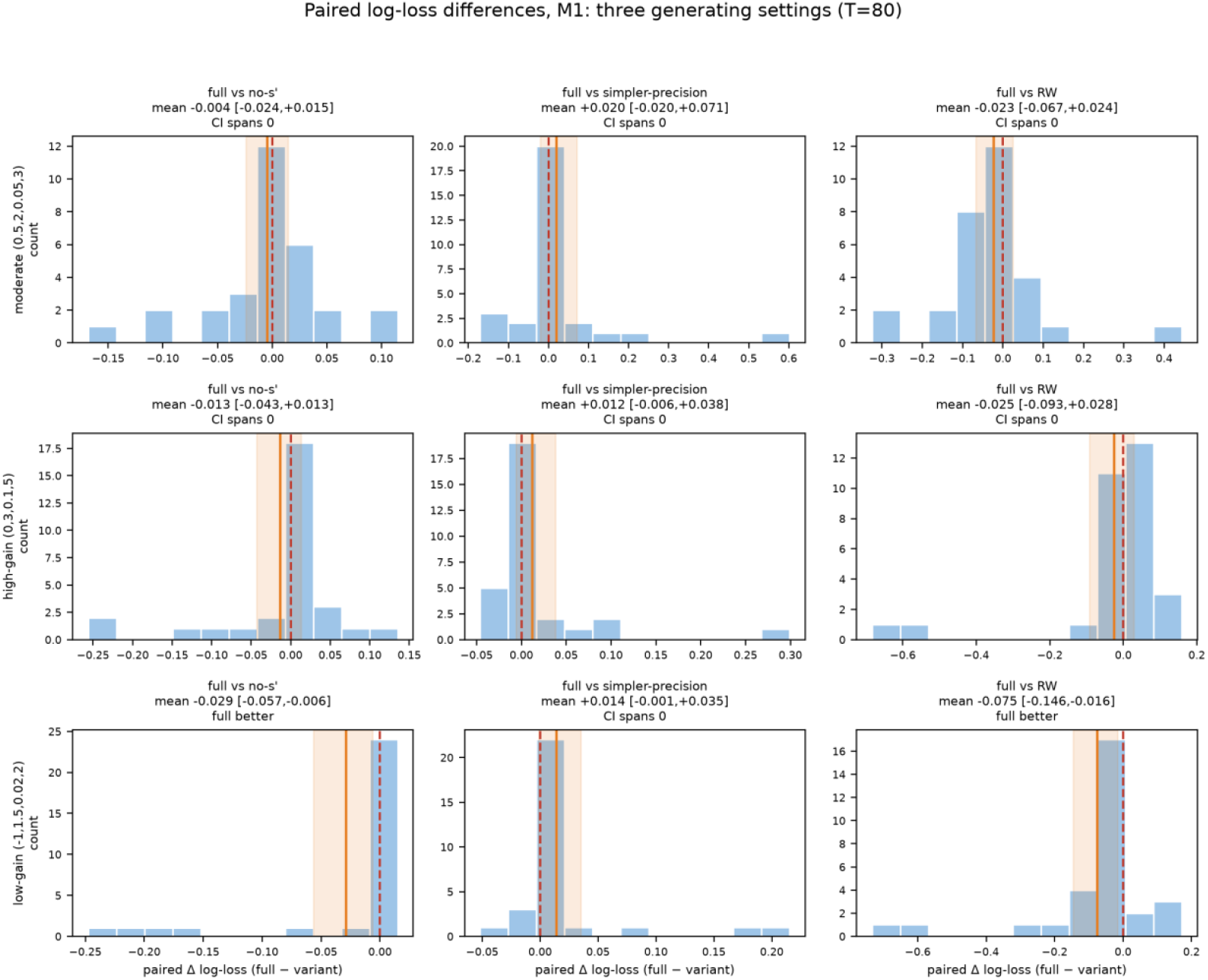
Experiment M1 (T=80 fixed): per-dataset paired log-loss differences (full − variant) for three settings (moderate, high-gain, low-gain; rows) and three comparisons (columns: mean-without-s′, simpler-precision, RW). Rugs show n=30 dataset differences; red dashed line = 0; orange line = mean; orange band = paired-bootstrap 95% CI (10,000 resamples). Panels are significant only if the CI excludes 0. Only low-gain full vs mean-without-s′ and full vs RW exclude 0.

**S4 Fig.**
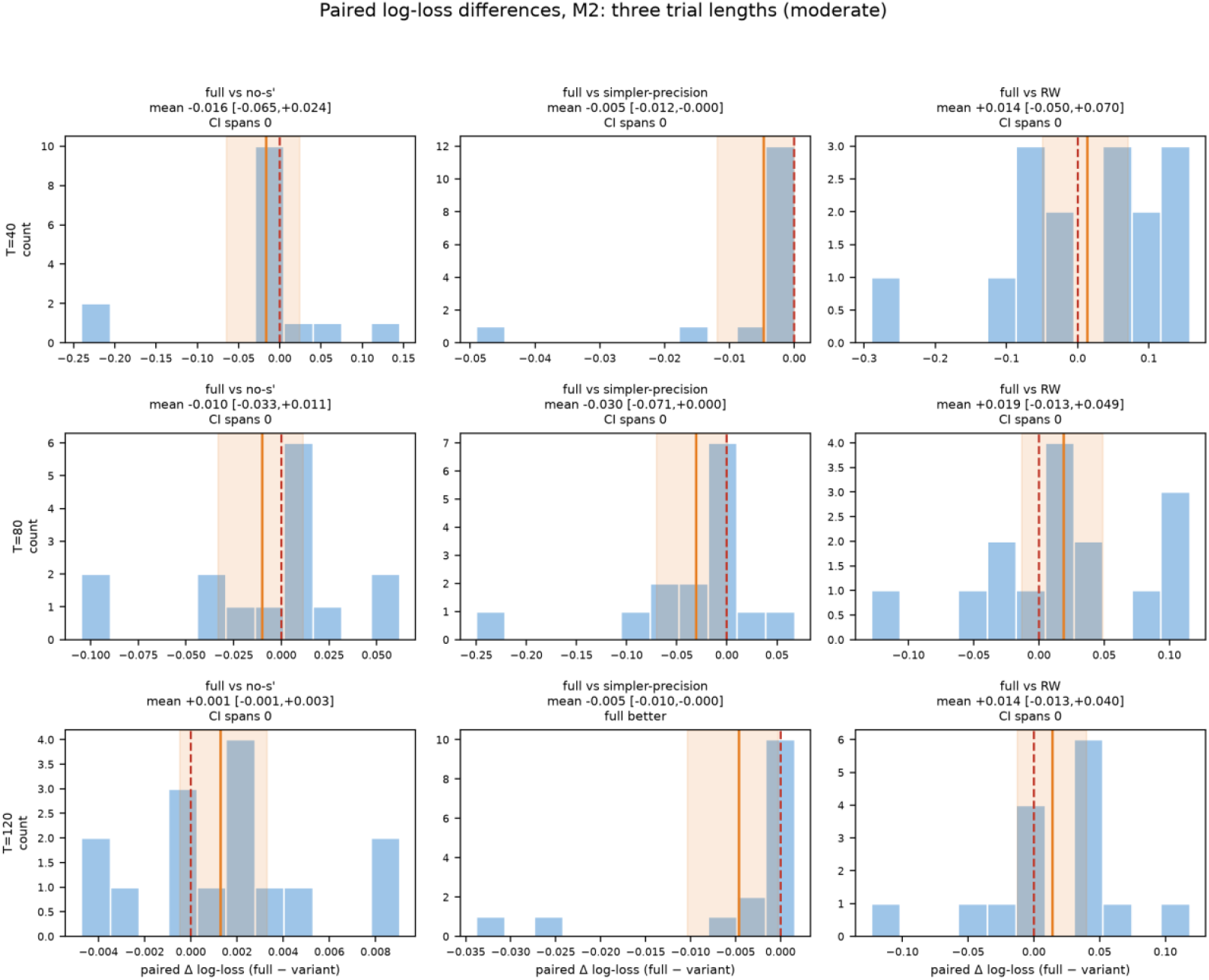
Experiment M2 (moderate): per-dataset paired log-loss differences for trial lengths T ∈ {40,80,120} (rows) and the same three comparisons (columns), laid out as in Fig. S3 (n=15 datasets/panel; red 0 line; orange mean with bootstrap-95% band). All nine normal-approximation intervals include zero. Two bootstrap upper bounds are slightly negative only at the 4th–5th decimal (−0.00002 at T=40; -0.00032 at T=120 for simpler-precision), and the T=80 bound (+0.00032) changes sign across resampling seeds. Overall, no panel shows a consistent or meaningful advantage for the full update rule over this trial-length range.

